# Systems genetics analyses in Diversity Outbred mice inform human bone mineral density GWAS and identify *Qsox1* as a novel determinant of bone strength

**DOI:** 10.1101/2020.06.24.169839

**Authors:** Basel M. Al-Barghouthi, Larry D. Mesner, Gina M. Calabrese, Daniel Brooks, Steven M. Tommasini, Mary L. Bouxsein, Mark C. Horowitz, Clifford J. Rosen, Kevin Nguyen, Samuel Haddox, Emily A. Farber, Suna Onengut-Gumuscu, Daniel Pomp, Charles R. Farber

## Abstract

Genome-wide association studies (GWASs) for osteoporotic traits have identified over 1000 associations; however, their impact has been limited by the difficulties of causal gene identification and a strict focus on bone mineral density (BMD). Here, we used Diversity Outbred (DO) mice to directly address these limitations by performing the first systems genetics analysis of 55 complex skeletal phenotypes. We applied a network approach to cortical bone RNA-seq data to discover 72 genes likely to be causal for human BMD GWAS associations, including the novel genes *SERTAD4* and *GLT8D2*. We also performed GWAS in the DO for a wide-range of bone traits and identified *Qsox1* as a novel gene influencing cortical bone accrual and bone strength. Our results provide a new perspective on the genetics of osteoporosis and highlight the ability of the mouse to inform human genetics.

## INTRODUCTION

Osteoporosis is a condition of low bone strength and an increased risk of fracture ^1^. It is also one of the most prevalent diseases in the U.S., affecting over 10 million individuals^2^. Over the last decade, efforts to dissect the genetic basis of osteoporosis using genome-wide association studies (GWASs) of bone mineral density (BMD) have been tremendously successful, identifying over 1000 independent associations ^3–5^. These data have the potential to revolutionize our understanding of bone biology and discovery of novel therapeutic targets ^6, 7^; however, progress to date has been limited.

One of the main limitations of human BMD GWAS is the difficulty identifying causal genes. This is largely due to the fact that most associations implicate non-coding variation presumably influencing BMD by altering gene regulation ^5^. For other diseases, the use of molecular “-omics” data (*e.g.*, transcriptomic, epigenomic, etc.) in conjunction with systems genetics approaches (*e.g.*, identification of expression quantitative trait loci (eQTL) and network-based approaches) has successfully informed gene discovery ^8, 9^. However, few “-omics” datasets exist on bone or bone cells in large human cohorts (*e.g*, bone or bone cells were not part of the Gene Tissue Expression (GTEx) project ^10^), limiting the use of systems genetics approaches to inform BMD GWAS ^11^.

A second limitation is that all large-scale GWASs have focused exclusively on BMD ^3–5^. BMD is a clinically relevant predictor of osteoporotic fracture; however, it explains only part of the variance in bone strength ^12–15^. Imaging modalities and bone biopsies can be used to collect data on other bone traits such as trabecular microarchitecture and bone formation rates; however, it will be difficult to apply these techniques “at scale” (N=>100K). Additionally, many aspects of bone, including biomechanical properties, cannot be measured *in vivo*. These limitations have hampered the dissection of the genetics of osteoporosis and highlight the need for resources and approaches that address the challenges faced by human studies.

The Diversity Outbred (DO) is a highly engineered mouse population derived from eight genetically diverse inbred founders (A/J, C57BL/6J, 129S1/SvImJ, NOD/ShiLtJ, NZO/HILtJ, CAST/EiJ, PWK/PhJ, and WSB/EiJ) ^16^. The DO has been randomly mated for over 30 generations and, as a result, it enables high-resolution genetic mapping and relatively efficient identification of causal genes ^17, 18^. As an outbred stock, the DO also more closely approximates the highly heterozygous genomes of a human population. These attributes, coupled with the ability to perform detailed and in-depth characterization of bone traits and generate molecular data on bone, position the DO as a platform to assist in addressing the limitations of human studies described above.

Here, we created a resource for the systems genetics of bone strength consisting of information on 55 bone traits from over 600 DO mice, and RNA-seq data from marrow-epleted cortical bone in 192 DO mice. We demonstrated the utility of this resource in two ways. First, we applied a network approach to the bone transcriptomics data in the DO and identified 72 genes that were bone-associated nodes in Bayesian networks, and their human homologs were located in BMD GWAS loci and regulated by colocalizing eQTL in human tissues. Of the 72, 21 were not previously known to influence bone. The further investigation of two of the 21 novel genes, *SERTAD4* and *GLT8D2*, revealed that they were likely causal and influenced BMD via a role in osteoblasts. Second, we performed GWASs in the DO for 55 complex traits associated with bone strength; identifying 28 QTL. By integrating QTL and bone eQTL data in the DO, we identified *Qsox1* as the gene responsible for a QTL on Chromosome (Chr.) 1 influencing cortical bone accrual along the medial-lateral femoral axis and femoral strength. These data highlight the power of the DO mouse resource to complement and inform human genetic studies of osteoporosis.

## RESULTS

### Development of a resource for the systems genetics of bone strength

An overview of the resource is presented in **Figure 1**. We measured 55 complex skeletal phenotypes in a cohort of DO mice (N=619; 314 males, 305 females; breeding generations 23-33) at 12 weeks of age. We also generated RNA-seq data from marrow-depleted femoral diaphyseal bone from a randomly chosen subset of the 619 phenotyped mice (N=192; 96/sex). All 619 mice were genotyped using the GigaMUGA^19^ array (∼110K SNPs) and these data were used to reconstruct the genome-wide haplotype structures of each mouse. As expected, the genomes of DO mice consisted of approximately 12.5% from each of the eight

**Figure 1.**
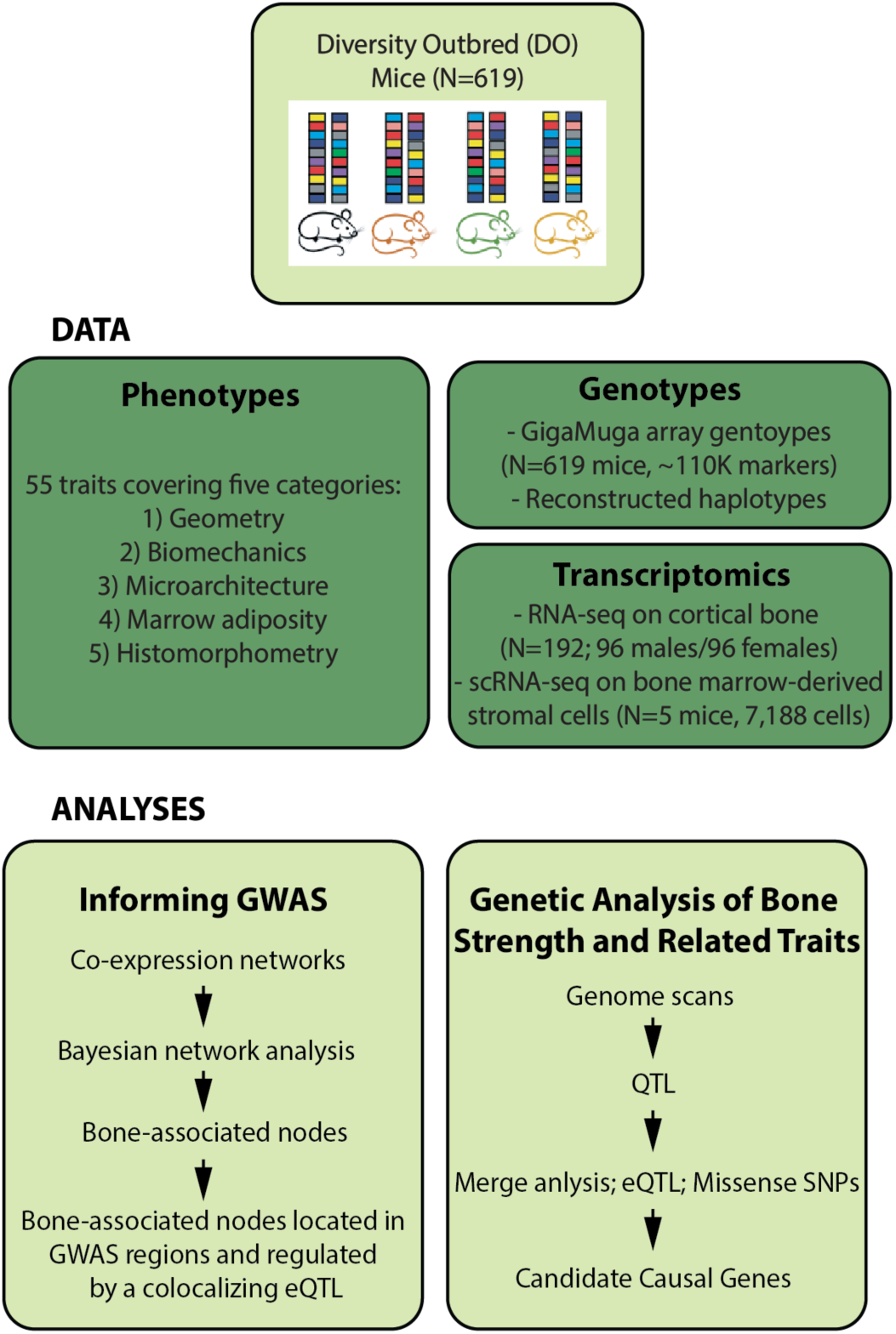
Resource overview. An overview of the resource including data generated and analyses performed.

DO founders (**Figure 2A**).

**Figure 2.**
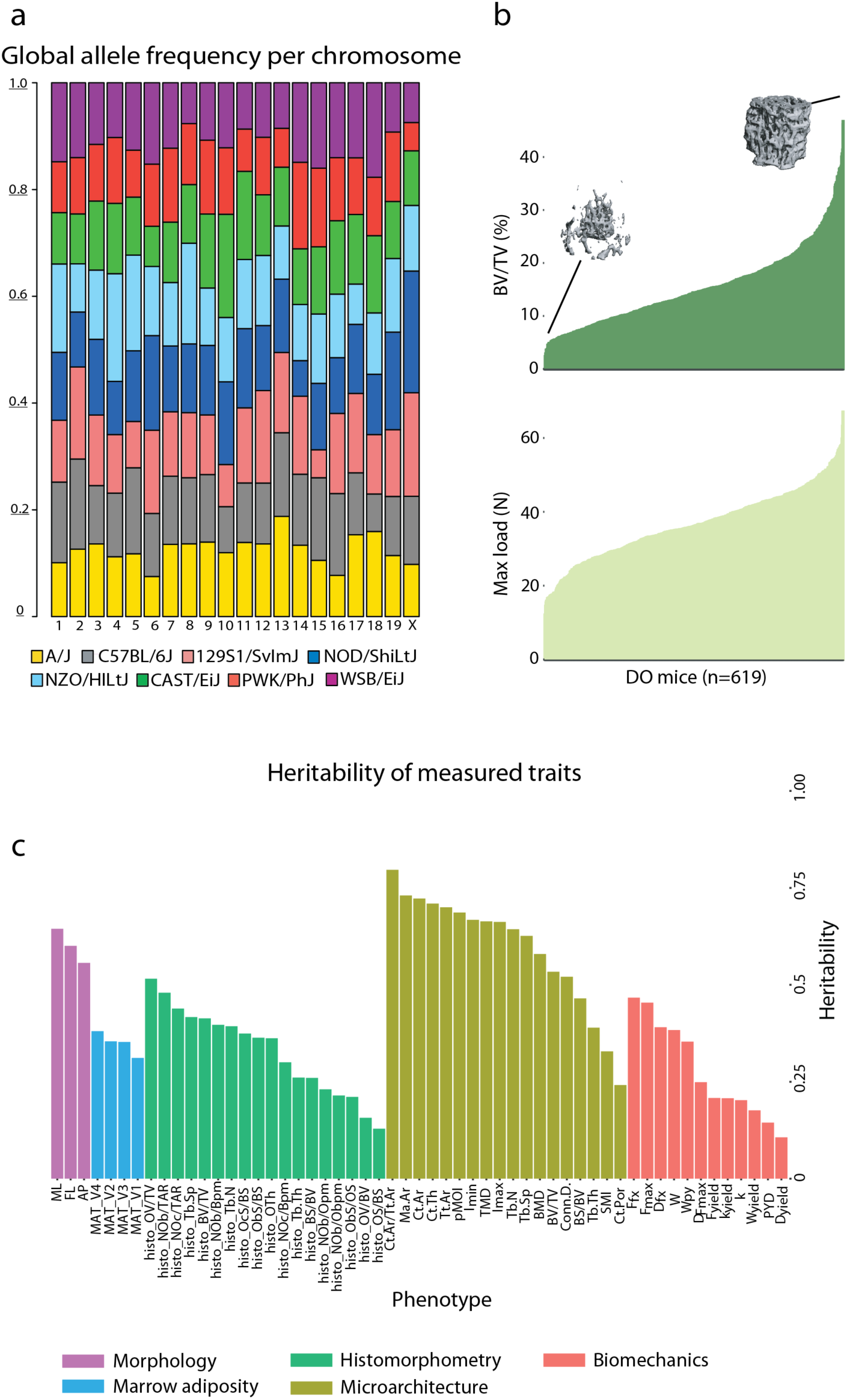
Characterization of the experimental cohort. **a**) Allele frequency per chromosome, across the DO mouse cohort. **b**) Bone volume fraction and max load across the DO cohort. Insets are microCT images representing low and high bone volume fraction. **c**) Heritability of each trait. Phenotypes are colored by phenotypic category.

The collection of phenotypes included measures of bone morphology, microarchitecture, and biomechanics of the femur, along with tibial histomorphometry and marrow adiposity (**Supplemental Tables 1 and 2)**. Our data included quantification of femoral strength as well as many clinically relevant predictors of strength and fracture risk (*e.g.*, trabecular and cortical microarchitecture). Traits in all categories (except tibial marrow adipose tissue (MAT)) were significantly (P_adj_<0.05) correlated with femoral strength (**Supplemental Table 3**). Additionally, all traits exhibited substantial variation across the DO cohort. For example, we observed a 30.8-fold variation (the highest measurement was 30.8 times greater than the lowest measurement) in trabecular bone volume fraction (BV/TV) of the distal femur and 5.6-fold variation in femoral strength (**Figure 2B)**. After adjusting for covariates (age, DO generation, sex, and body weight) all traits had non-zero heritabilities (h^2^) (**Figure 2C**). Correlations between traits in the DO were consistent with expected relationships observed in previous mouse and human studies (**Supplemental Table 4)** ^20–23^.

**Table 1.**
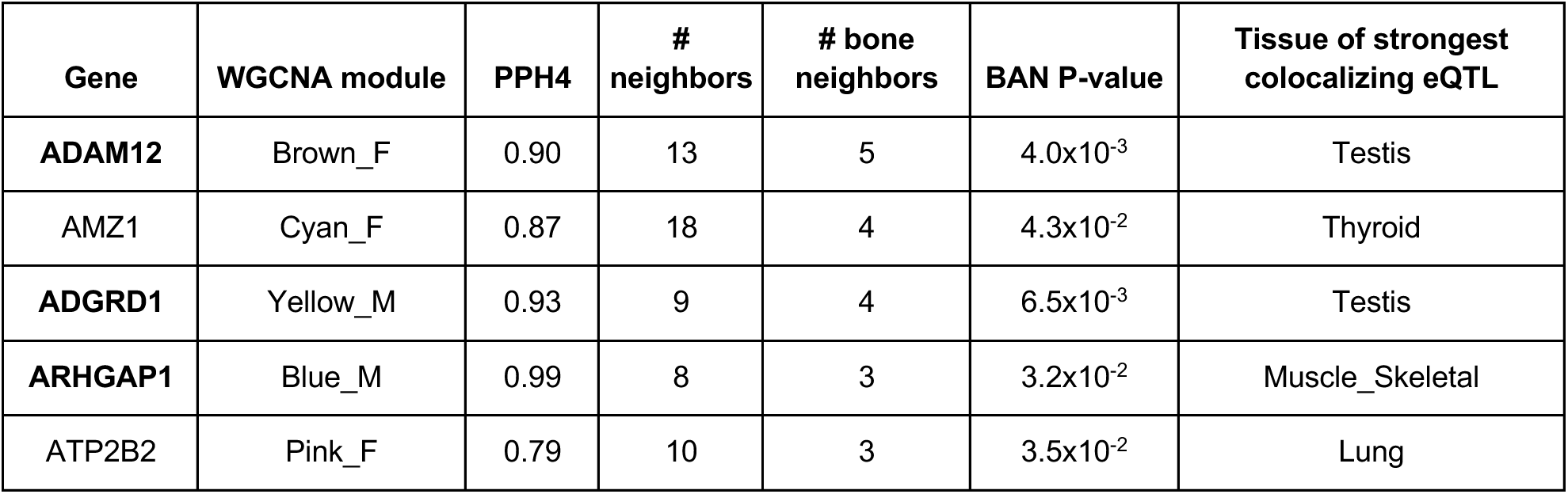
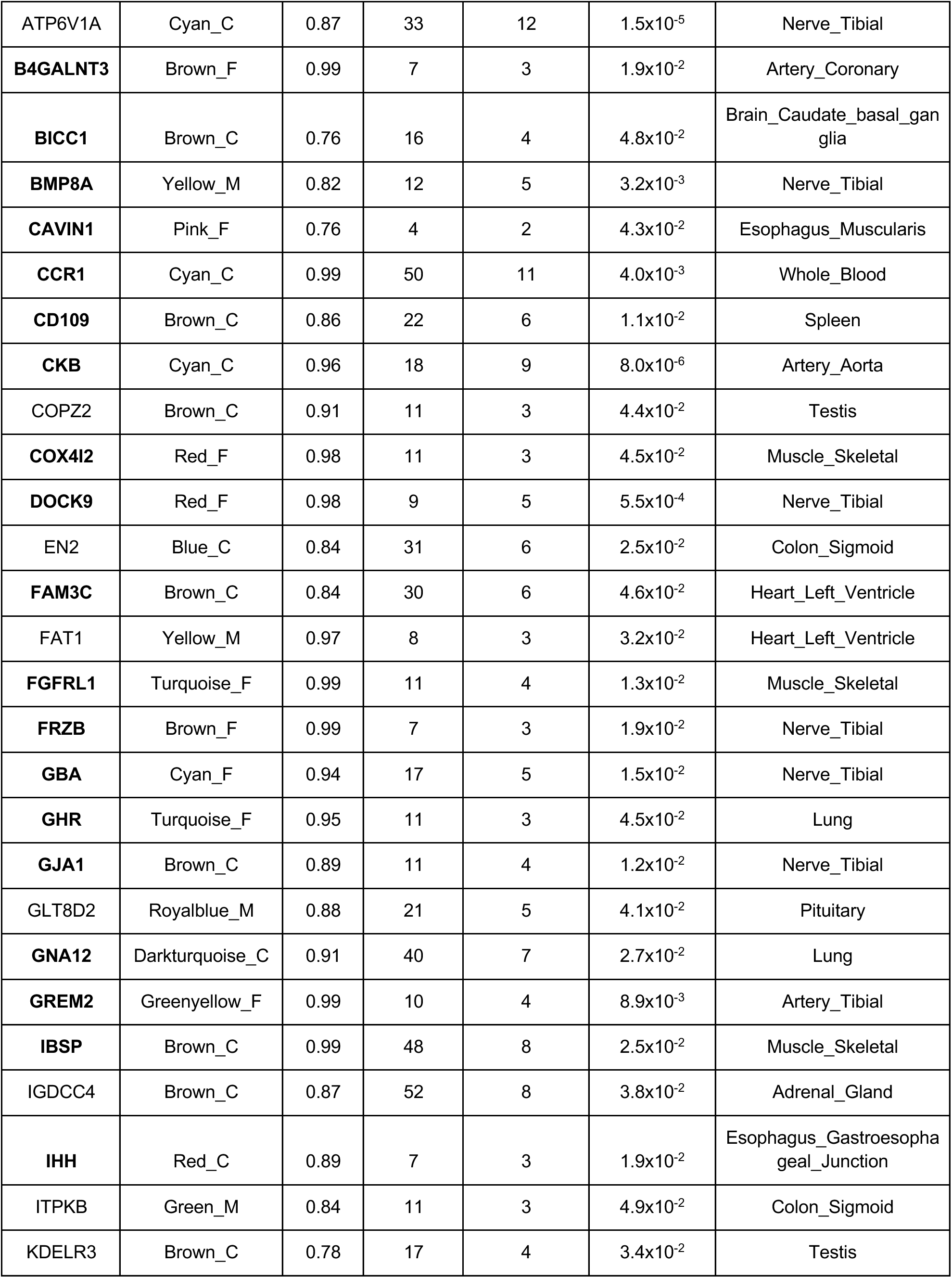
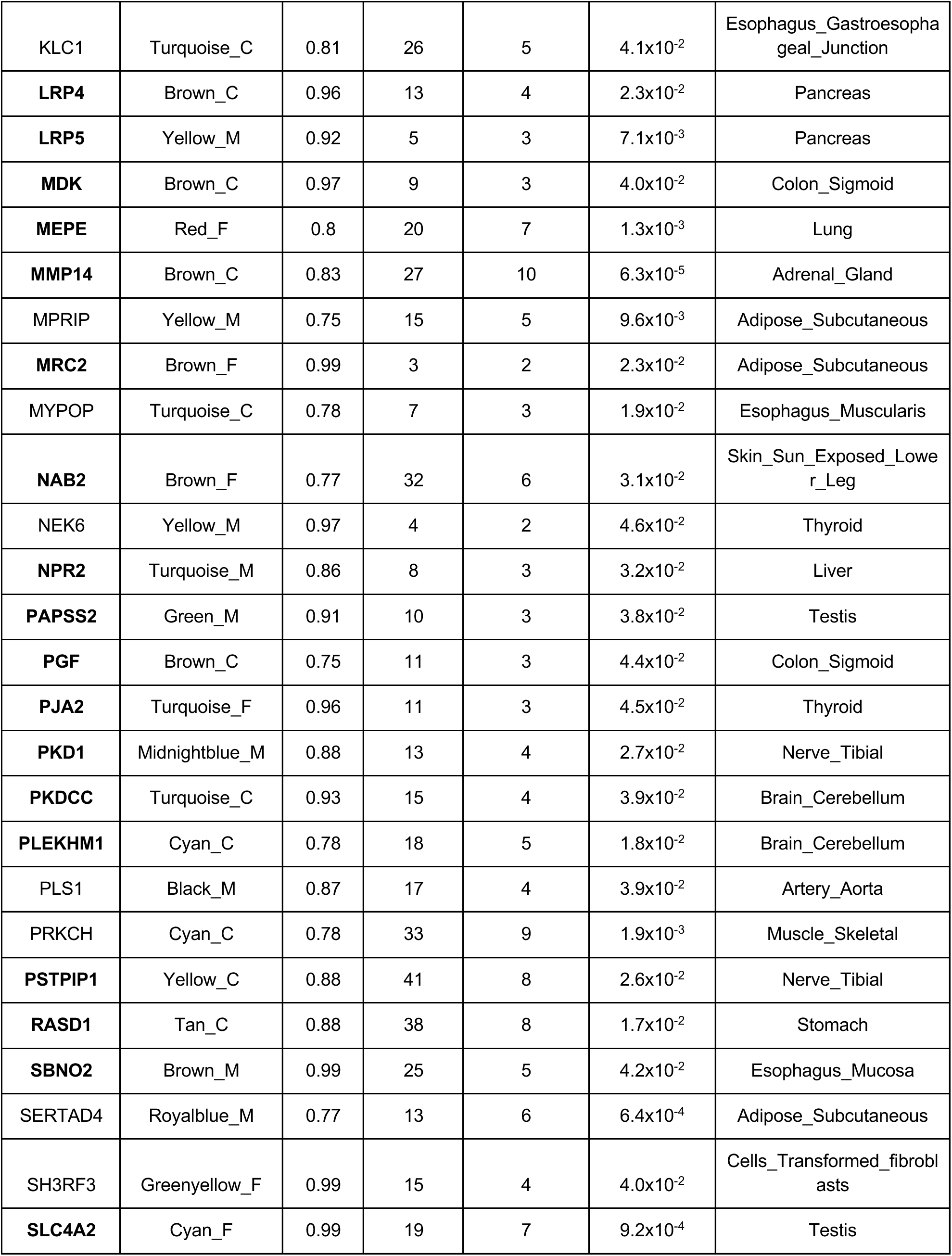
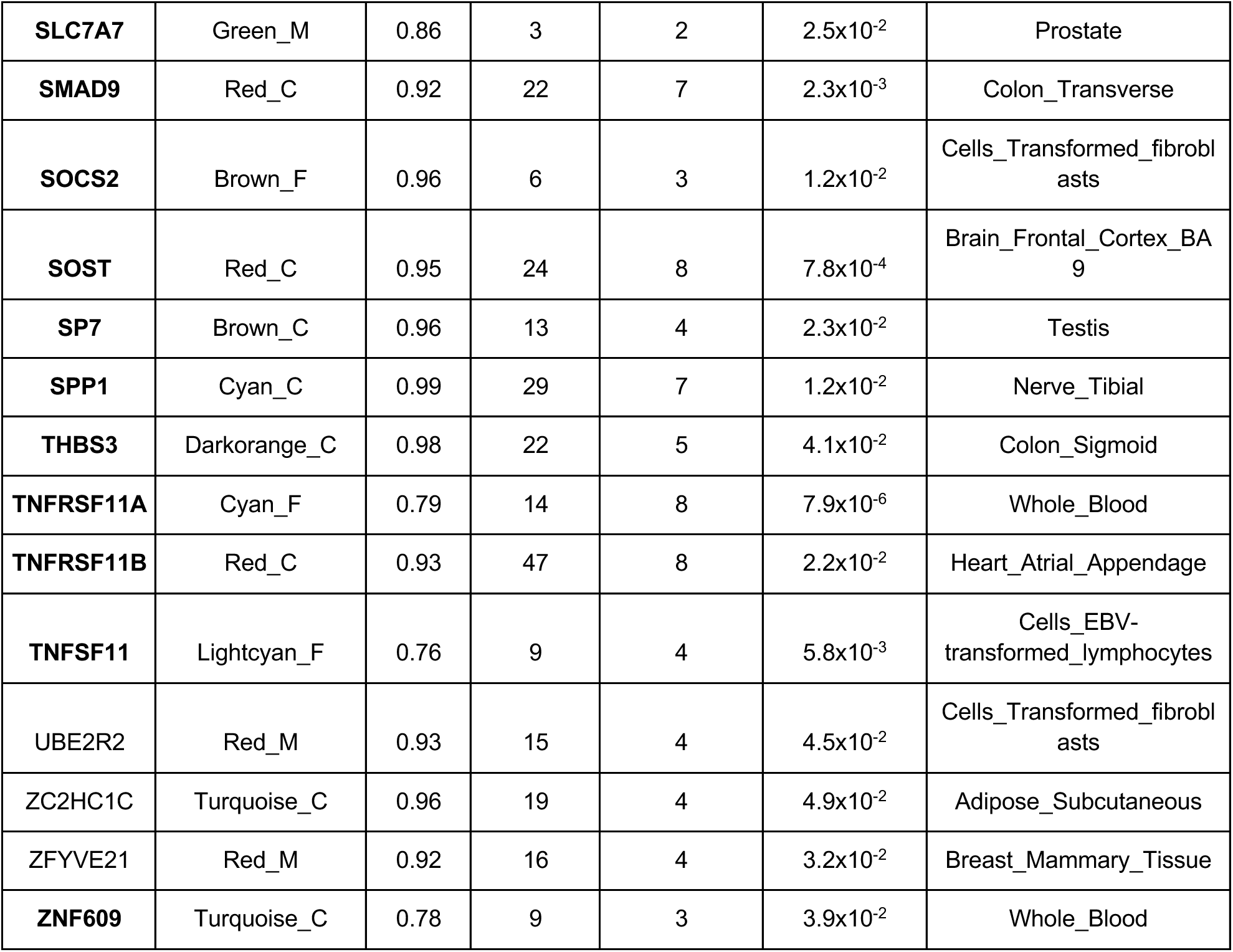
Homologous human BANs with colocalizing eQTL. Genes in bold are known bone genes.

**Table 2.**
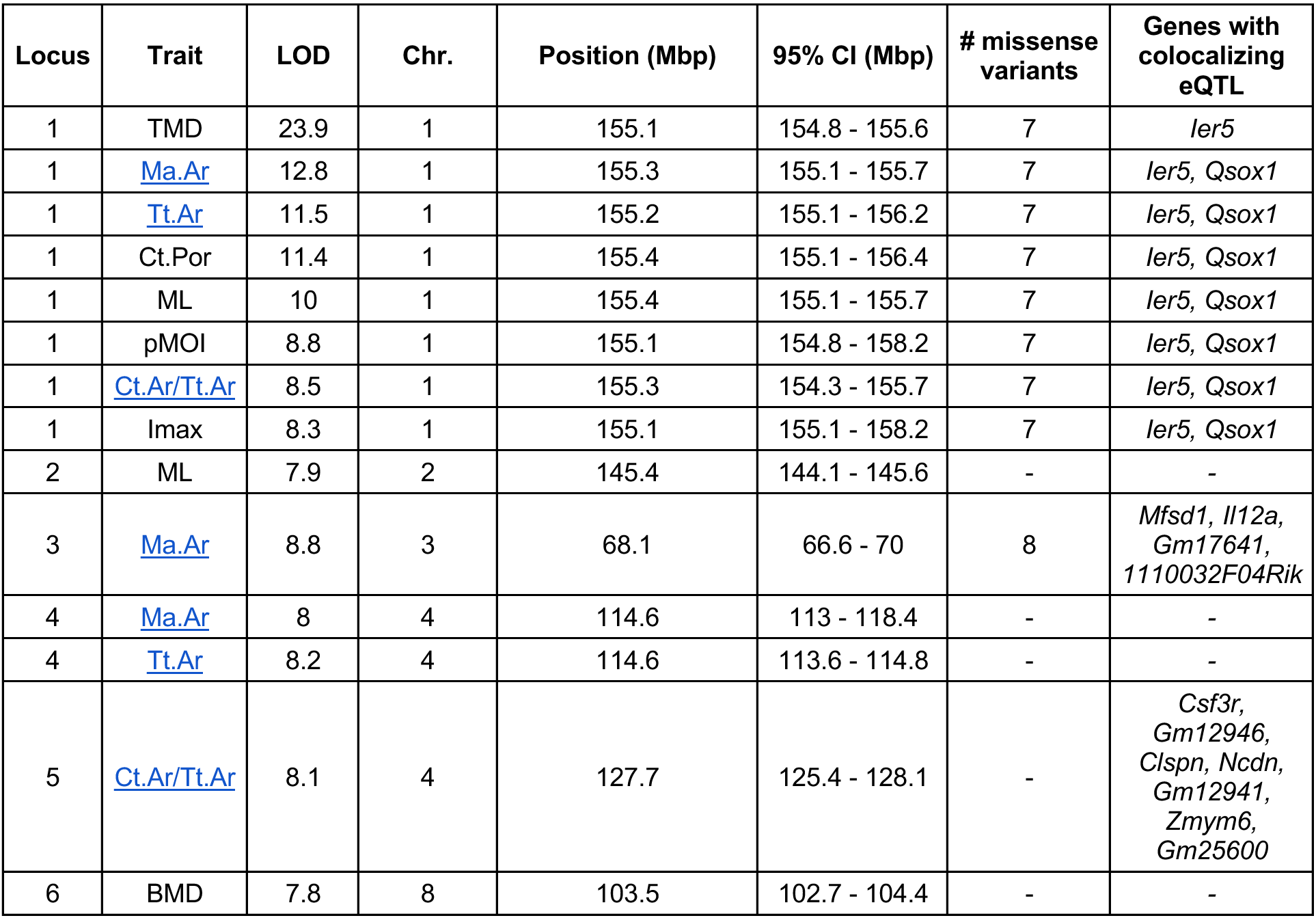
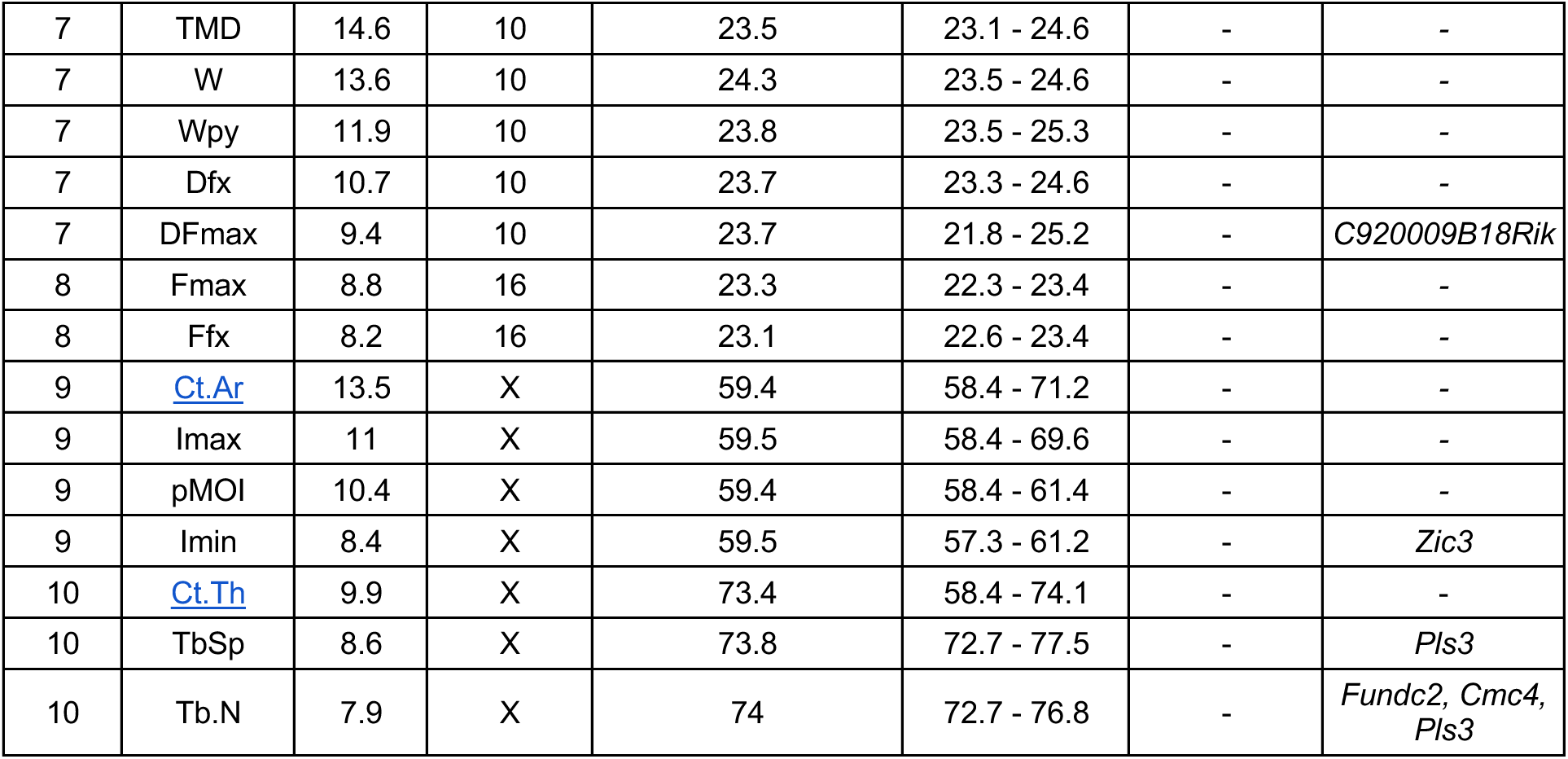
QTL identified for complex skeletal traits in the DO.

In addition to standard RNA-seq quality control procedures (**Methods**), we also assessed RNA-seq quality by principal components analysis (PCA) and as expected, we identified effects of sex (PC3) and batch (**Supplemental Figure 1**), which we then corrected for prior to downstream analyses. Importantly, our PCA analysis did not identify any outliers in the bulk RNA-seq data Furthermore, we performed differential expression analyses between sexes and between individuals with high versus low bone strength (**Supplemental Tables 5 and 6**). As expected, the most significantly differentially expressed genes based on sex were located on the X chromosome. We identified 83 significantly (FDR<0.05) differentially-expressed transcripts in the analysis of low and high bone strength. Many were genes, such as *Ahsg* ^24^ and *Arg1* ^25^, which have previously been implicated in the regulation of bone traits.

### Identification of bone-associated nodes

We wanted to address the challenge of identifying causal genes from BMD GWAS data, using the DO resource described above. To do so, we employed a network-based approach similar to one we have used in prior studies ^26, 27^ (**Figure 3**). First, we partitioned genes into groups based on co-expression by applying weighted gene co-expression network analysis (WGCNA) to the DO cortical bone RNA-seq data ^28^. We generated three WGCNA networks; sex-combined, male, and female. The three networks contained a total of 124 modules (**Supplemental Table 7**). A Gene Ontology (GO) analysis revealed that nearly all modules were enriched for genes involved in specific biological processes, including modules enriched for processes specific to bone cells (osteoblasts or osteoclasts) (**Supplemental Table 8**).

**Figure 3.**
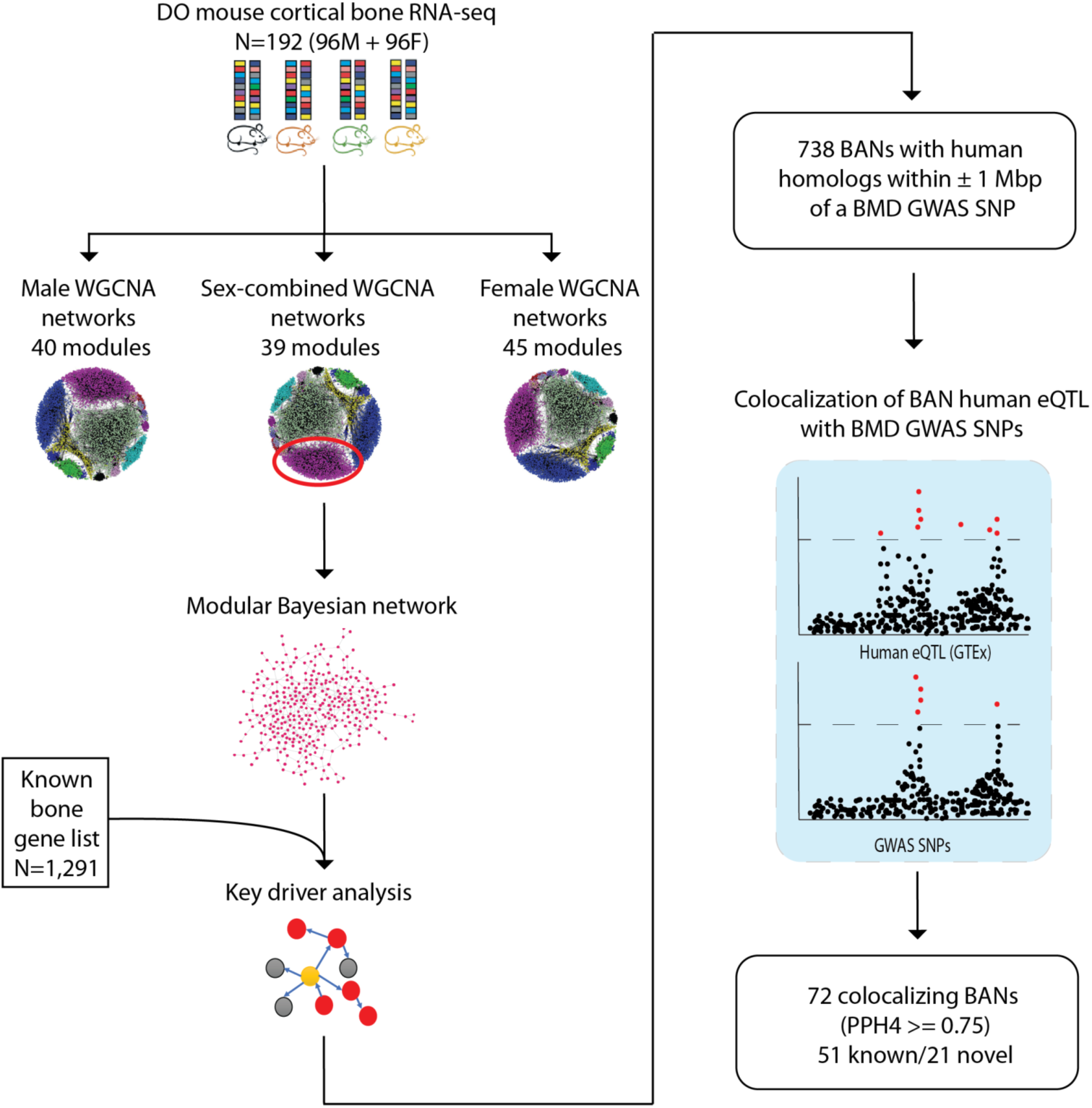
Overview of the network approach used to identify genes potentially responsible for BMD GWAS loci. Three WGCNA networks (124 total modules) were constructed from RNA-seq data on cortical bone in the DO (N=192). A Bayesian network was then learned for each module. We performed key driver analysis on each Bayesian network to identify bone-associated nodes (BANs), by identifying nodes (genes) that were more connected to more known bone genes than was expected by chance. We colocalized GTEx human eQTL for each BAN with GWAS BMD SNPs to identify potentially causal genes at BMD GWAS loci.

We next sought to infer causal interactions between genes in each module, and then use this information to identify genes likely involved in regulatory processes relevant to bone and the regulation of BMD. To do so, we generated Bayesian networks for each co-expression module, allowing us to model directed gene-gene relationships based on conditional independence. Bayesian networks allowed us to model causal links between co-expressed (and likely co-regulated) genes.

We hypothesized that key genes involved in bone regulatory processes would play central roles in bone networks and, thus, be more highly connected in the Bayesian networks. In order to test this hypothesis, we generated a list of genes implicated in processes known to impact bone or bone cells (“known bone gene” list (N=1,291); **Supplemental Table 9**; see **Methods)**. The GWAS loci referenced in this study were enriched in human homologs of genes in the “known bone list”, relative to the set of protein-coding genes in the genome (OR=1.35, P=1.45^-7^). Across the three network sets (combined, male and female), we found that genes with putative roles in bone regulatory processes were more highly connected than all other genes (P=2.6 x 10^-5^, P=5.2 x 10^-3^, and P=5.5 x 10^-7^ for combined, male, and female network sets, respectively), indicating the structure of the Bayesian networks was not random with respect to connectivity.

To discover genes potentially responsible for GWAS associations, we identified bone-associated nodes (BANs). BANs were defined as genes connected in our Bayesian networks with more genes in the “known bone gene” list than would be expected by chance ^29–32^. The analysis identified 1,465 genes with evidence (P_nominal_<0.05) of being a BAN (*i.e.,* sharing network connections with genes known to participate in a bone “regulatory” process) (**Supplemental Table 10**).

### Using BANs to inform human BMD GWAS

We reasoned that the BAN list was enriched for causal BMD GWAS genes. In fact, of the 1,465 BANs, 1,251 had human homologs and 738 of those were within 1 Mbp of one of the 1,161 BMD GWAS lead SNPs identified in ^3^ and ^5^. This represents an enrichment of BANs within GWAS loci (+/- 1Mbp of GWAS SNP), relative to the number of protein-coding genes within GWAS loci (OR=1.28, P=1.97 x 10^-5^).

However, a gene being a BAN is likely not strong evidence, by itself, that a particular gene is causal for a BMD GWAS association. Therefore, to provide additional evidence connecting BMD-associated variants to the regulation of BANs, we identified local eQTL for each BAN homolog in 48 human non-bone samples using the Gene Tissue Expression (GTEx) project ^10, 33, 34^. Our rationale for using GTEx was that while these data do not include information on bone tissues or bone cells, a high degree of local eQTL sharing has been observed between GTEx tissues ^10, 35^. This suggests that a colocalizing eQTL in a non-bone tissue may represent either a non-bone autonomous causal effect or may reflect the actions of a shared eQTL that is active in bone and shared across non-bone tissues. We then tested each eQTL for colocalization (*i.e.*, probability the eQTL and GWAS association share a common causal variant) with their respective BMD GWAS association ^3,5^. Of the 738 BANs located in proximity of a BMD GWAS locus, 72 had colocalizing eQTL (PPH4≥0.75, **Supplemental Table 11,** see **Methods**) in at least one GTEx tissue (**Table 1**). Of these, 51 (70.8%) were putative regulators of bone traits (based on comparing to the known bone gene list (N=40) and a literature search for genes influencing bone cell function (N=11)), highlighting the ability of the approach to recover known biology. Based on overlap with the known bone gene list, this represents a highly significant enrichment of known bone genes in the list of BANs with colocalizing eQTL relative to the number of known bone genes in the list of GWAS-proximal BANs (OR=2.53, P=1.65 x 10^-4^). Our approach identified genes such as *SP7* (Osterix) ^36^, *SOST* ^37, 38^, and *LRP5* ^39–41^, which play central roles in osteoblast-mediated bone formation. Genes essential to osteoclast activity, such as *TNFSF11* (RANKL) ^42–45^, *TNFRSF11A* (RANK) ^46, 47^, and *SLC4A2* ^48^ were also identified. Twenty-one (29.2%) genes were not previously implicated in the regulation of bone traits.

One of the advantages of the network approach is the ability to identify potentially causal genes and provide insight into how they may impact BMD based on their module membership and network connections. For example, the “cyan” module in the female network (cyan_F) harbored many of the known BANs that influence BMD through a role in osteoclasts (the GO term “osteoclast differentiation” was highly enriched P=2.8 x 10^-15^ in the cyan_F module) (**Supplemental Table 8**). Three of the twenty-one novel BANs with colocalizing eQTL (**Table 1**), *ATP6V1A*, *PRKCH* and *AMZ1*, were members of the cyan module in the female network. Based on their cyan module membership it is likely they play a role in osteoclasts. *ATP6V1A* is a subunit of the vacuolar ATPase V1 domain ^49^. The vacuolar ATPase plays a central role in the ability of osteoclasts to acidify matrix and resorb bone, though *ATP6V1A* itself (which encodes an individual subunit) has not been directly connected to the regulation of BMD ^49^. *PRKCH* encodes the eta isoform of protein kinase C and is highly expressed in osteoclasts ^50^. *AMZ1* is a zinc metalloprotease and is relatively highly expressed in osteoclasts, and is highly expressed in macrophages, which are osteoclast precursors ^50^.

Next, we focused on two of the novel BANs with colocalizing eQTL, *SERTAD4* (GTEx Adipose Subcutaneous; coloc PPH4=0.77; PPH4/PPH3=7.9) and *GLT8D2* (GTEx Pituitary; coloc PPH4=0.88; PPH4/PPH3=13.4). Both genes were members of the royalblue module in the male network (royalblue_M). The function of *SERTAD4* (SERTA domain-containing protein 4) is unclear, though proteins with SERTA domains have been linked to cell cycle progression and chromatin remodeling ^51^. *GLT8D2* (glycosyltransferase 8 domain containing 2) is a glycosyltransferase linked to nonalcoholic fatty liver disease ^52^. In the DO, the eigengene of the royalblue_M module was significantly correlated with several traits, including trabecular number (Tb.N; rho=-0.26; P=9.5 x 10^-3^) and separation (Tb.Sp; rho=0.27; P=7.1 x 10^-3^), among others (**Supplemental Table 12**). The royalblue_M module was enriched for genes involved in processes relevant to osteoblasts such as “extracellular matrix” (P=8.4 x 10^-19^), “endochondral bone growth” (P=5.7 x 10^-4^), “ossification” (P=8.9 x 10^-4^) and “negative regulation of osteoblast differentiation” (P=0.04) (**Supplemental Table 8**). Additionally, *Sertad4* and *Glt8d2* were connected, in their local (3-step) Bayesian networks, to well-known regulators of osteoblast/osteocyte biology (such as *Wnt16* ^53^, *Postn* ^54, 55^, and *Col12a1* ^56^ for *Sertad4* and *Pappa2* ^57^, *Pax1* ^57, 58^, and *Tnn* ^59^ for *Glt8d2*) (**Figure 4A and 4B**). *Sertad4* and *Glt8d2* were strongly expressed in calvarial osteoblasts with expression increasing (P<2.2 x 10^-16^ and P=6.4 x 10^-10^, respectively) throughout the course of differentiation (**Figure 4C**). To further investigate their expression in osteoblasts, we generated single-cell RNA-seq (scRNA-seq) data on mouse bone marrow-derived stromal cells exposed to osteogenic differentiation media *in vitro* from our mouse cohort (N=5 mice (4 females, 1 male), 7,188 cells, **Supplemental Table 13**, **Supplemental Figure 2**). Clusters of cell-types were grouped into mesenchymal progenitors, preadipocytes/adipocytes, osteoblasts, osteocytes, and non-osteogenic cells based on the expression of genes defining each cell-type (**Supplemental Table 14**). *Sertad4* was expressed across multiple cell-types, with its highest expression in a specific cluster (cluster 9) of mesenchymal progenitor cells and lower levels of expression in osteocytes (cluster 10) (**Figure 4D, Supplemental Figure 3**). *Glt8d2* was expressed in a relatively small number of cells in both progenitor and mature osteoblast populations (**Figure 4D, Supplemental Figure 3**).

**Figure 4.**
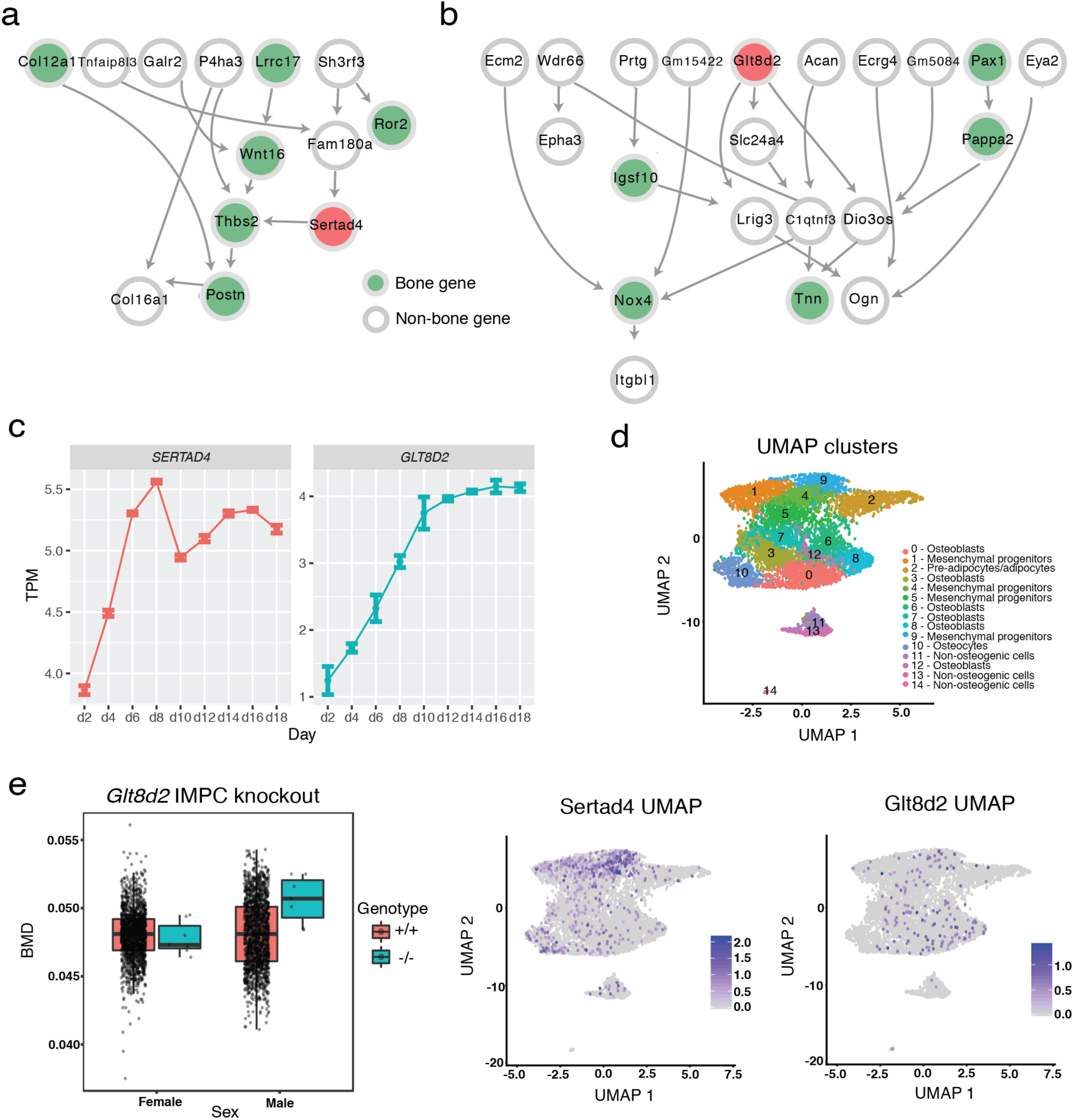
Identifying *SERTAD4* and *GLT8D2* as putative regulators of BMD. **a**) Local 3-step neighborhood around *Sertad4*. Known bone genes highlighted in green. *Sertad4* highlighted in red. **b**) Local 3-step neighborhood around *Glt8d2*. Known bone genes highlighted in green. *Glt8d2* highlighted in red. **c**) Expression of *Sertad4* and *Glt8d2* in calvarial osteoblasts. Error bars represent the standard error of the mean. **d**) Single cell RNA-seq expression data. Each point represents a cell. The top panel shows UMAP clusters and their corresponding cell-type. The bottom two panels show the expression of *Sertad4* and *Glt8d2.* **e**) Bone mineral density in *Glt8d2* knockout mice from the IMPC. The center line is the median, and the whiskers represent 1.5 x IQR.

Finally, we analyzed data from the International Mouse Phenotyping Consortium (IMPC) for *Glt8d2* ^60^. After controlling for body weight, there was a significant (P=1.5 x 10^-3^) increase in BMD in male *Glt8d2*^-/-^ and no effect (P=0.88) in female *Glt8d2*^-/-^ mice (sex interaction P= 6.9 x 10^-3^) **(Figure 4E)**. These data were consistent with the direction of effect predicted by the human *GLT8D2* eQTL and eBMD GWAS locus where the effect allele of the lead eBMD SNP (rs2722176) was associated with increased *GLT8D2* expression and decreased BMD. Together, these data suggest that *SERTAD4* and *GLT8D2* are causal for their respective BMD GWAS associations and they likely impact BMD through a role in modulating osteoblast-centric processes.

### Identification of QTLs for strength-related traits in the DO

The other key limitation of human genetic studies of osteoporosis has been the strict focus on BMD, though many other aspects of bone influence its strength. To directly address this limitation using the DO, we performed GWAS for 55 complex skeletal traits. This analysis identified 28 genome-wide significant (permutation-derived P<0.05) QTLs for 20 traits mapping to 10 different loci (defined as QTL with peaks within a 1.5 Mbp interval) (**Table 2 and Supplemental Figure 4**). These data are presented interactively in a web-based tool (http://qtlviewer.uvadcos.io/). Of the 10 loci, four impacted a single trait (*e.g.,* medial-lateral femoral width (ML) QTL on Chr2@145.4Mbp), while the other six impacted more than one trait (*e.g.*, cortical bone morphology traits, cortical tissue mineral density (TMD), and cortical porosity (Ct.Por) QTL on Chr1@155Mbp). The 95% confidence intervals (CIs) for the 21 autosomal associations ranged from 615 Kbp to 5.4 Mbp with a median of 1.4 Mbp.

### Overlap with human BMD GWAS

We anticipated the genetic analysis of bone strength traits in DO mice would uncover novel biology not captured by human BMD GWAS. To evaluate this prediction, we identified overlaps between the 10 identified mouse loci and human BMD GWAS associations ^3,5^. Of the 10 mouse loci, the human syntenic regions (**Supplemental Table 15**) for six (60%) contained at least one independent GWAS association (**Supplemental Figure 5**). We calculated the number expected by chance by randomly selecting 10 human regions (of the same size) 1000 times, followed by identifying overlaps. Six overlaps corresponded to the 57^th^ percentile of the null distribution.

### Identification of potentially causal genes

For each locus, we defined the causal gene search space as the widest confidence interval given all QTL start and end positions ±250 Kbp. We then used a previously described approach, merge analysis, to fine-map QTL and identify likely causal genes (**Figure 5**) ^61^. Merge analysis was performed by imputing all known variants from the genome sequences of the eight founders onto haplotype reconstructions for each DO mouse, and then performing single variant association tests. We focused on variants in the top 15% of each merge analysis as those are most likely to be causal ^61^.

**Figure 5.**
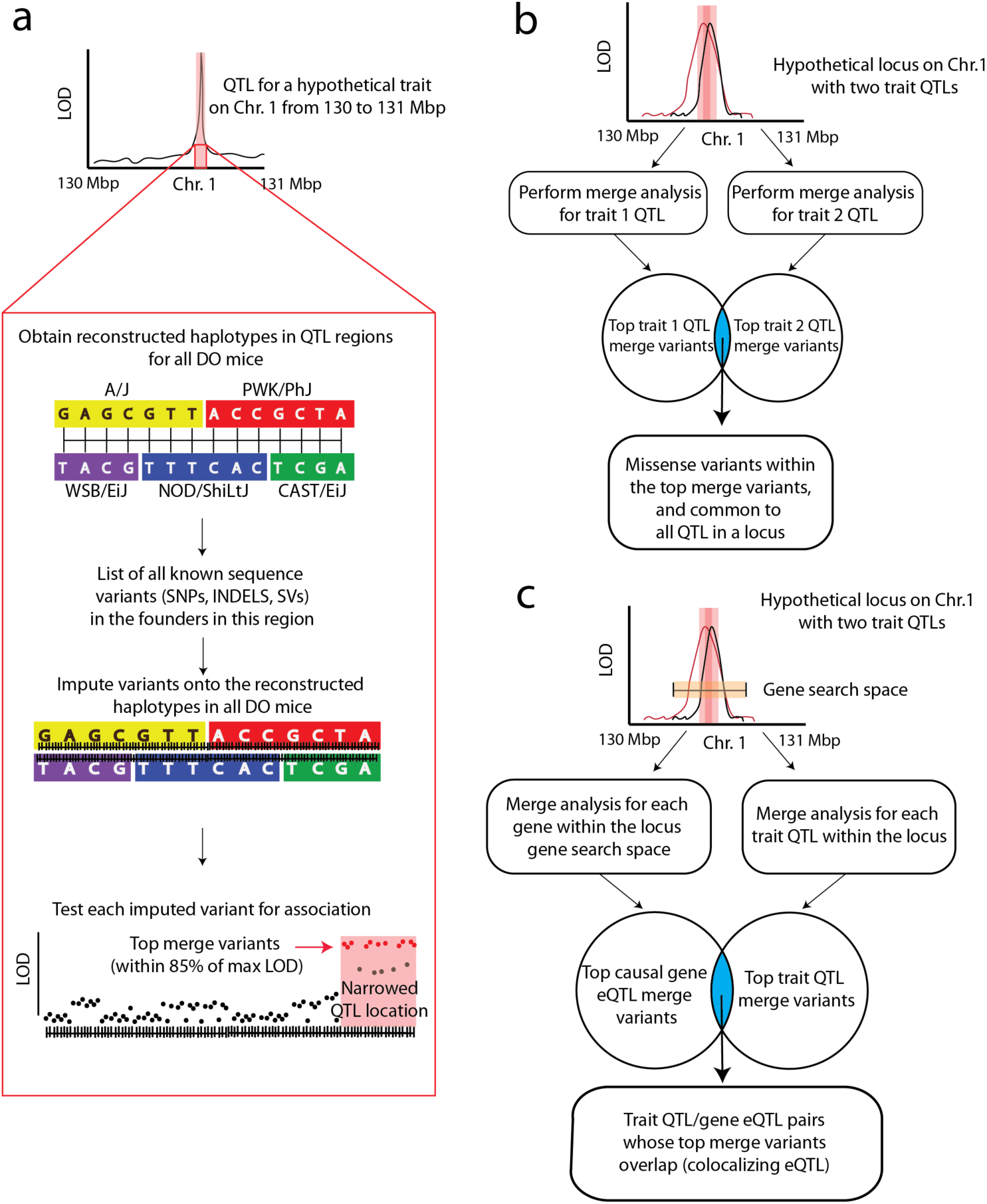
Overview of our approach to QTL fine-mapping. **a**) Overview of merge analysis. **b**) Overview of merge analysis as performed for the identification of missense variants. **c**) Overview of merge analysis as performed for the identification of colocalizing trait QTL/ gene eQTL within a locus. The pink columns around the QTL in each association plot represent the QTL 95% confidence intervals. The yellow box in panel (**c**) represents the gene search space for a locus, defined as the region within ± 250 Kbp around the outer boundaries of the 95% confidence intervals within a locus.

We next identified missense variants that were top merge analysis variants common to all QTL in a locus. We identified seven missense variants in locus 1, and eight missense variants in locus 3 (**Table 2**). Of the seven missense variants in locus 1, three (rs243472661, rs253446415, and rs33686629) were predicted to be deleterious by SIFT. They are all variants in the uncharacterized protein coding gene *BC034090*. In locus 3, three (rs250291032, rs215406048 and rs30914256) were predicted to be deleterious by SIFT (**Supplemental Table 16**). These variants were located in myeloid leukemia factor 1 (*Mlf1*), Iqcj and Schip1 fusion protein (*Iqschfp*), and Retinoic acid receptor responder 1 (*Rarres1*), respectively.

We next used the cortical bone RNA-seq data to map 10,399 local eQTL in our DO mouse cohort (**Supplemental Table 17**). Of these, 174 local eQTL regulated genes located within bone trait QTL. To identify colocalizing eQTL, we identified trait QTL/eQTL pairs whose top merge analysis variants overlapped. This analysis identified 18 genes with colocalizing eQTL in 6 QTL loci (**Table 2**).

### Characterization of a QTL on Chromosome 1 influencing bone morphology

Locus 1 (Chr1) influenced cortical bone morphology (Ma.Ar, total cross sectional area (Tt.Ar), medial-lateral femoral width (ML), polar moment of inertia (pMOI), cortical bone area fraction (Ct.Ar/Tt.Ar), and maximum moment of inertia (I_max_)), tissue mineral density (TMD), and cortical porosity (Ct.Por) (**Figure 6A**). We focused on this locus due to its strong effect size and the identification of candidate genes (*Ier5*, *Qsox1,* and *BC034090*) (**Table 2**). Additionally, we had previously measured ML in an independent cohort of DO mice (N=577; 154 males/423 females) from earlier generations (generations G10 and G11) and a QTL scan of those data uncovered the presence of a similar QTL on Chr1 ^62^ (**Supplemental Figure 6, Methods**). The identification of this locus across two different DO cohorts (which differed in generations, diets, and ages) provided robust replication justifying further analysis.

**Figure 6.**
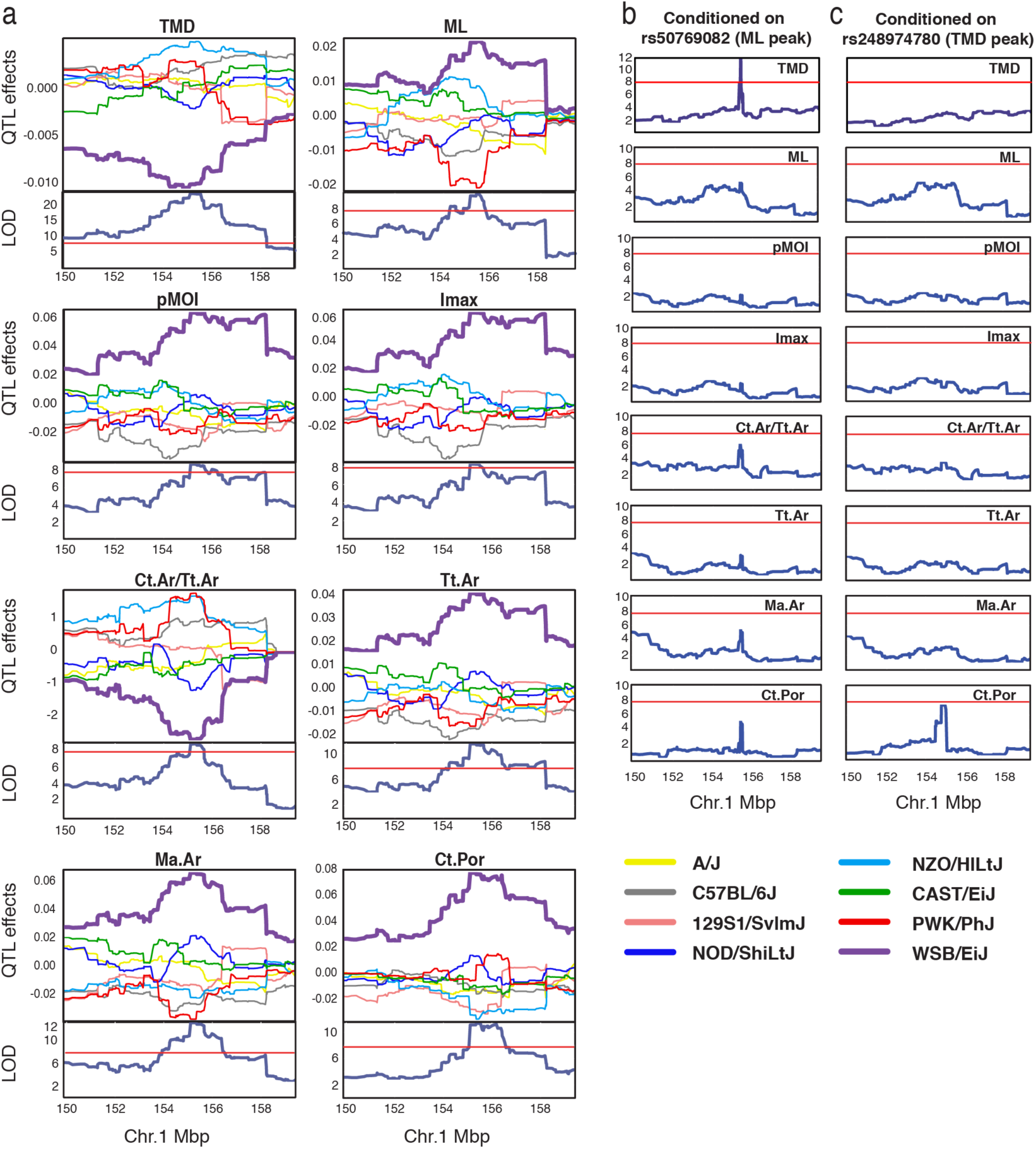
QTL (locus 1) on chromosome 1. **a**) For each plot, the top panel shows allele effects for the DO founders for each of the 8 QTL across an interval on chromosome 1 (Mbp, colors correspond to the founder allele in the legend). Bottom panels show each respective QTL scan. The red horizontal lines represent LOD score thresholds (Genome-wide P<=0.05). **b**) QTL scans across the same interval as panel (A), after conditioning on rs50769082. **c**) QTL scans after conditioning on rs248974780.

The traits mapping to this locus fell into two phenotypic groups, those influencing different aspects of cross-sectional size (e.g., ML and Tt.Ar) and TMD/cortical porosity. We suspected that locus 1 QTL underlying these two groups were distinct, and that QTL for traits within the same phenotypic group were linked. This hypothesis was further supported the observation that correlations among the size traits were strong and cross-sectional size traits were not correlated with TMD or porosity **(Supplemental Table 4)**. Therefore, we next tested if the locus affected all traits or was due to multiple linked QTL. The non-reference alleles of the top merge analysis variants for each QTL were private to WSB/EiJ. To test if these variants explained all QTL, we performed the same association scans for each trait, but included the genotype of the lead ML QTL variant (rs50769082; 155.46 Mbp; ML was used as a proxy for all the cortical morphology traits) as an additive covariate. This led to the ablation of all QTL except for TMD which remained significant (**Figure 6B**). We then repeated the analysis using the lead TMD QTL variant (rs248974780; 155.06 Mbp) as an additive covariate (**Figure 6C**). This led to the ablation of all QTLs. These results supported the presence of at least two loci both driven by WSB/EiJ alleles, one influencing cortical bone morphology and Ct.Por and the other influencing TMD, as well as possibly influencing cortical bone morphology and Ct.Por.

### *Qsox1* is responsible for the effect of locus 1 on cortical bone morphology

Given the importance of bone morphology to strength, we sought to focus on identifying the gene(s) underlying locus 1 and impacting cortical bone morphology. We re-evaluated candidate genes in light of the evidence for two distinct QTL. Immediate Early Response 5 (*Ier5)* and quiescin sulfhydryl oxidase 1 (*Qsox1*) were identified as candidates based on the DO mouse eQTL analysis and *BC034090* as a candidate based on missense variants (**Table 2**). Interestingly, *Ier5* and *Qsox1* eQTL colocalized with all QTL, except the TMD QTL, where only *Ier5* colocalized, providing additional support for two distinct loci (**Table 2 and Figure 7A**). We cannot exclude the involvement of the missense variants in *BC034090*; however, without direct evidence that they impacted *BC034090* function, we put more emphasis on the eQTL. As a result, based on its colocalizing eQTL and known biological function (see below), we predicted that *Qsox1* was at least partially responsible for locus 1.

**Figure 7.**
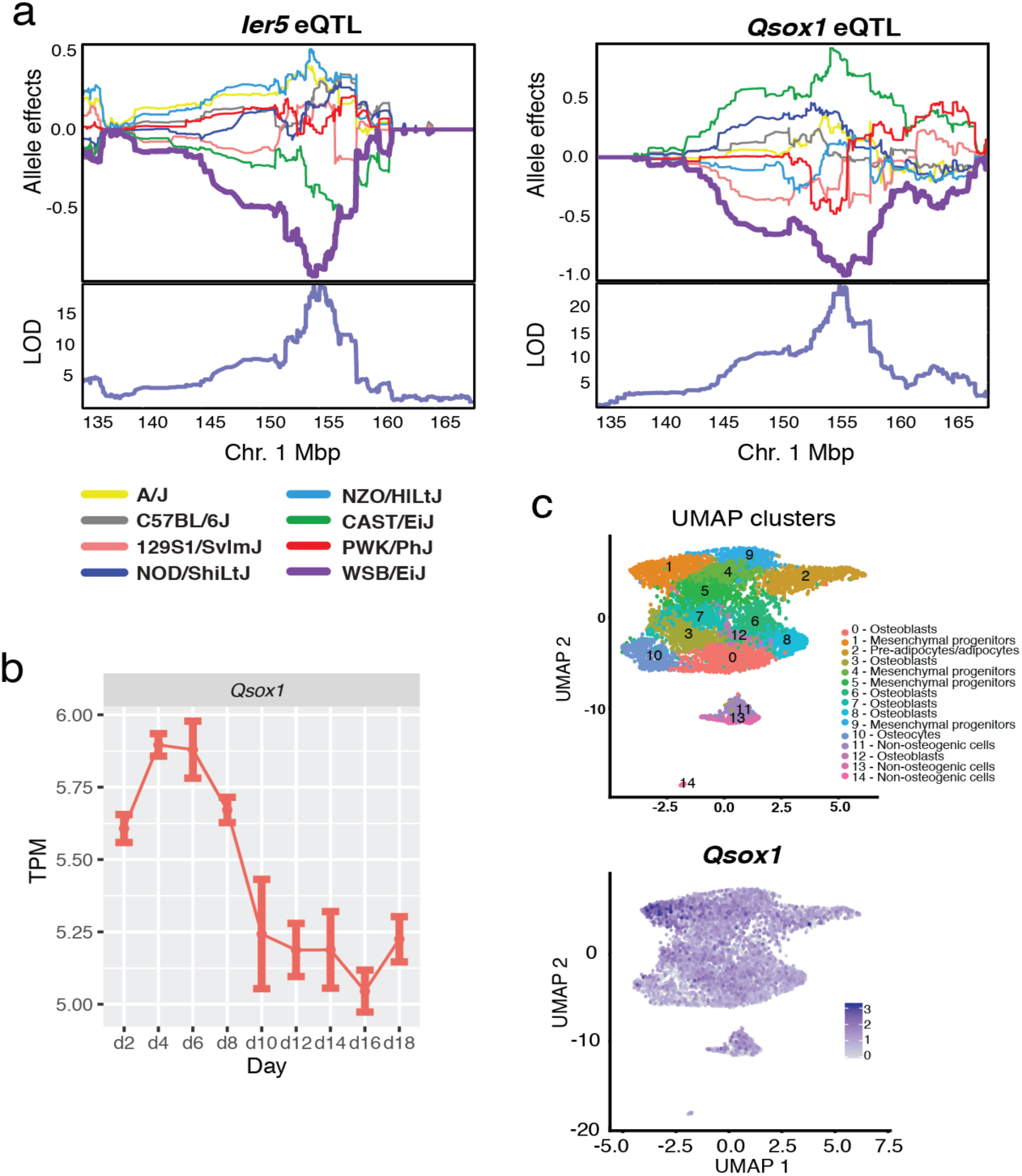
Characterization of *Qsox1*. **a**) The top panel shows allele effects for the DO founders for *Ier5* and *Qsox1* expression an interval on chromosome 1 (Mbp, colors correspond to the founder allele in the legend). Y-axis units are best linear unbiased predictors (BLUPs). Bottom panels show each respective QTL scan. LOD score threshold for autosomal eQTL is 10.89 (alpha=0.05). **b**) *Qsox1* expression in calvarial osteoblasts. Error bars represent the standard error of the mean. **c**) Single cell RNA-seq expression data. Each point represents a cell. The top panel shows UMAP clusters and their corresponding cell-type. The bottom panel shows the expression of *Qsox1*.

QSOX1 is the only known secreted catalyst of disulfide bond formation and a regulator of extracellular matrix integrity ^63^. It has not been previously linked to skeletal development. We found that *Qsox1* was highly expressed in calvarial osteoblasts and its expression decreased (P=6.4 x 10^-6^) during differentiation (**Figure 7B**). In scRNA-seq on bone marrow-derived stromal cells exposed to osteogenic differentiation media *in vitro*, we observed *Qsox1* expression in all osteogenic cells with its highest expression seen in a cluster of mesenchymal progenitors defined by genes involved in skeletal development such as *Grem2*, *Lmna*, and *Prrx2* (cluster 1) (**Supplemental Table 18 and Figure 7C**). Additionally, in the DO cortical bone RNA-seq data, *Qsox1* was highly co-expressed with many key regulators of skeletal development and osteoblast activity (*e.g.*, *Runx2*; rho=0.48, P=<2.2 x 10^-16^, *Lrp5*; rho=0.41, P=6.2 x 10^-9^).

To directly test the role of *Qsox1*, we used CRISPR/Cas9 to generate *Qsox1* mutant mice. We generated five different mutant lines harboring unique mutations, including two 1-bp frameshifts, a 171-bp in-frame deletion of the QSOX1 catalytic domain, and two large deletions (756 bp and 1347 bp) spanning most of the entire first exon of *Qsox1* (**Figure 8A, Supplemental Tables 19 and 20**). All five mutations abolished QSOX1 activity in serum (**Figure 8B**). Given the uniform lack of QSOX1 activity, we combined phenotypic data from all lines to evaluate the effect of QSOX1 deficiency on bone. We hypothesized based on the genetic and DO mouse eQTL data, that QSOX1 deficiency would increase all traits mapping to locus 1, except TMD. Consistent with this prediction, ML was increased overall (P=1.8 x10^-9^), and in male (P=5.6×10^-7^) and female (P=3.5×10^-3^) mice as a function of *Qsox1* mutant genotype (**Figure 8C**). Also consistent with the genetic data, we observed no difference in other gross morphological traits including anterior-posterior femoral width (AP) (P=0.31) (**Figure 8D**) and femoral length (FL) (P=0.64) (**Figure 8E**). We next focused on male *Qsox1*^+/+^ and *Qsox1^-^*^/-^ mice and used microCT to measure other bone parameters. We observed increased pMOI (P=0.02) (**Figure 8F**), Imax (P=0.009) (**Figure 8G**), and Ct.Ar/Tt.Ar (P=0.031) (**Figure 8H**). Total area (Tt.Ar) (**Figure 8I**) was increased, but the difference was only suggestive (P=0.08). Marrow area (Ma.Ar, P=0.93) was not different (**Figure 8J**). We observed no change in TMD (P=0.40) (**Figure 8K**). We also observed no difference in cortical porosity (Ct.Por) (P=0.24) (**Figure 8L**).

**Figure 8.**
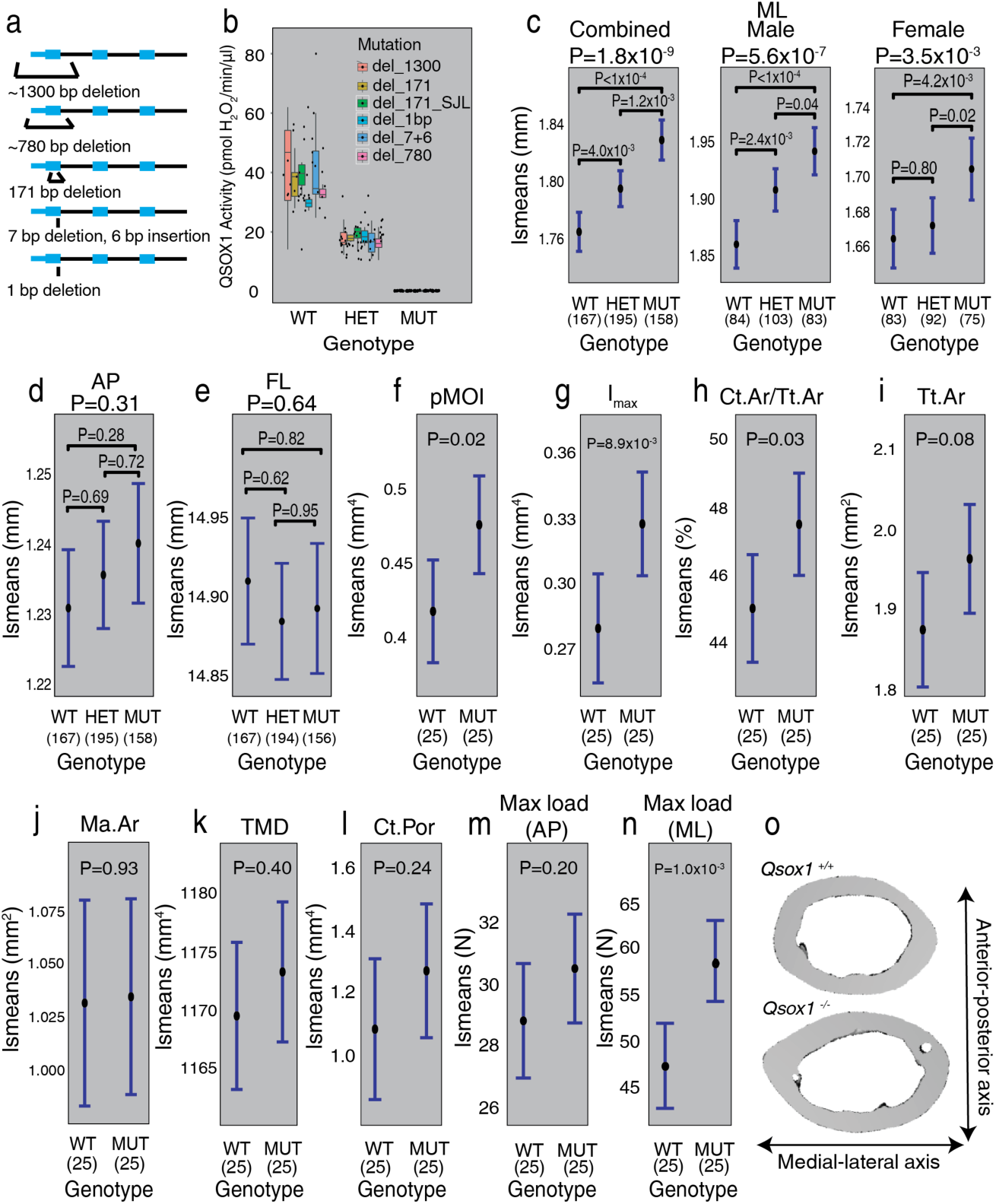
*Qsox1* is responsible for several chromosome 1 QTL. **a**) Representative image of the *Qsox1* knockout mutations. **b**) QSOX1 activity assay in serum. Data is grouped by mouse genotype. Boxplot whiskers represent 1.5 x IQR, and the center line is the median. **c-e**) Caliper-based measurements of femoral morphology in *Qsox1* mutant mice. P-values above plots are ANOVA P-values for the genotype term, while P-values in the plots are contrast P-values, adjusted for multiple comparisons. **f-l**) microCT measurements of chromosome 1 QTL phenotypes in *Qsox1* knockout mice. **m**) Bone strength (max load, F_max_) in the AP orientation, measured via four-point bending. **n**) Bone strength (max load, F_max_) in the ML orientation, measured via four-point bending. **o**) Representative microCT images of the effect of *Qsox1* on bone size. All error bars represent confidence intervals at a 95% confidence level.

Given the strength of locus 1 on bone morphology and its association with biomechanical strength, we were surprised the locus did not impact femoral strength. Typically, in four-point bending assays, the force is applied along the AP axis. We replicated this in femurs from *Qsox1*^+/+^ and *Qsox1^-^*^/-^ mice and saw no significant impact on strength (P=0.20) (**Figure 8M**). However, when we tested femurs by applying the force along the ML axis, we observed a significant increase in strength in *Qsox1^-^*^/-^ femurs (P=1.0 x 10^-3^) (**Figure 8N**). Overall, these data demonstrate that absence of QSOX1 activity leads to increased cortical bone accrual specifically along the ML axis (**Figure 8O**).

## Discussion

Human GWASs for BMD have identified over 1000 loci. However, progress in causal gene discovery has been slow and BMD explains only part of the variance in bone strength and the risk of fracture ^7^. The goal of this study was to demonstrate that systems genetics in DO mice can help address these limitations. Towards this goal, we used cortical bone RNA-seq data in the DO and a network-based approach to identify 72 genes likely causal for BMD GWAS loci. Twenty-one of the 72 were novel. We provide further evidence supporting the causality of two of these genes, *SERTAD4* and *GLT8D2*. Furthermore, GWAS in the DO identified 28 QTLs for a wide-range of strength associated traits. From these data, *Qsox1* was identified as a novel genetic determinant of cortical bone mass and strength. These data highlight the power of systems genetics in the DO and demonstrate the utility of mouse genetics to inform human GWAS and bone biology.

To inform BMD GWAS, we generated Bayesian networks for cortical bone and used them to identify BANs. Our analysis was similar to “key driver” analyses ^29–31^ where the focus has often been on identifying genes with strong evidence (P_adj_<0.05) of playing central roles in networks. In contrast, we used BAN analysis as a way to rank genes based on the likelihood (P_nominal_ ≤ 0.05) that they are involved in a biological process important to bone (based on network connections to genes known to play a role in bone biology). We then identified genes most likely to be responsible for BMD GWAS associations by identifying BANs regulated by human eQTL that colocalize with BMD GWAS loci. Together, a gene being both a BAN in a GWAS locus and having a colocalizing eQTL is strong support of causality. This is supported by the observation that ∼71% of the 72 BANs with colocalizing eQTL were putative regulators of bone traits, based on a literature review and overlap with the “known bone gene” list.

One advantage of our network approach was the ability to not only identify causal genes, but use network information to predict the cell-type through which these genes are likely acting. We demonstrate this idea by investigating the two novel BANs with colocalizing human eQTL from the royalblue_M module. The royalblue_M module was enriched in genes involved in bone formation and ossification, suggesting the module as a whole and its individual members were involved in osteoblast-driven processes. This prediction was supported by the role of genes in osteoblasts that were directly connected to *Sertad4* and *Glt8d2*, the expression of the two genes in osteoblasts, and for *Glt8d2*, its regulation of BMD *in vivo*. Little is known regarding the specific biological processes that are likely impacted by *Sertad4* and *Glt8d2* in osteoblasts; however, it will be possible to utilize this information in future experiments designed to investigate their specific molecular functions. For example, *Sertad4* was connected to *Wnt16*, *Ror2*, and *Postn* all of which play roles in various aspects of osteoblast/osteocyte function. Wnt signaling is a major driver of osteoblast-mediated bone formation and skeletal development ^64^. Interestingly, *Wnt16* and *Ror2* play central roles in canonical (*Wnt16*) and non-canonical (*Ror2* in the *Wnt5a*/*Ror2* pathway) Wnt signaling ^65^ and have been shown to physically interact in chondrocytes ^66^. *Postn* has also been shown to influence Wnt signaling ^66, 67^. These data suggest a possible role for *Sertad4* in Wnt signaling.

Despite their clinical importance, we know little about the genetics of bone traits other than BMD. Here, we set out to address this knowledge gap. Using the DO, we identified 28 QTL for a wide-range of complex bone traits. The QTL were mapped at high-resolution, most had 95% CIs < 1 Mbp ^18^. This precision, coupled with merge and eQTL analyses in DO mice, allowed us to identify a small number of candidate genes for many loci. Overlap of existing human BMD GWAS association and mouse loci was no more than what would be expected by chance, suggesting that our approach has highlighted biological processes impacting bone that are independent of those with the largest effects on BMD. This new knowledge has the potential to lead to novel pathways which could be targeted therapeutically to increase bone strength. Future studies extending the work presented here will lead to the identification of additional genes and further our understanding of genetics of a broad range of complex skeletal traits.

Using multiple approaches, we identified *Qsox1* as responsible for at least part of the effect of the locus on Chr. 1 impacting bone morphology. We use the term “at least part” because it is clear that the Chr. 1 locus is complex. Using ML width as a proxy for all the bone morphology traits mapping to Chr. 1, the replacement of a single WSB/EiJ allele was associated with an increase in ML of 0.064 mm. Based on this, if *Qsox1* was fully responsible for the Chr. 1 locus we would expect at least an ML increase of 0.128 mm in *Qsox1* knockout mice; however, the observed difference was 0.064 mm (50% of the expected difference). This could be due to differences in the effect of *Qsox1* deletion in the DO compared to the SJL x B6 background of the *Qsox1* knockout or to additional QTL in Chr. 1 locus. The latter is supported by our identification of at least two QTL in the region. Further work will be needed to fully dissect this locus.

Disulfide bonds are critical to the structure and function of numerous proteins ^68^. Most disulfide bonds are formed in the endoplasmic reticulum ^69^; however, the discovery of QSOX1 demonstrated that disulfide bonds in proteins can be formed extracellularly ^63^. Ilani et al. ^63^ demonstrated that fibroblasts deficient in QSOX1 had a decrease in the number of disulfide bonds in matrix proteins. Moreover, the matrix formed by these cells was defective in supporting cell-matrix adhesion and lacked incorporation of the alpha-4 isoform of laminin. QSOX1 has also been associated with perturbation of the extracellular matrix in the context of cancer and tumor invasiveness ^70, 71^. It is unclear at this point how QSOX1 influences cortical bone mass; however, it likely involves modulation of the extracellular matrix.

In summary, we have used a systems genetics analysis in DO mice to inform human GWAS and identify novel genetic determinants for a wide-range of complex skeletal traits. Through the use of multiple synergistic approaches, we have expanded our understanding of the genetics of BMD and osteoporosis. This work has the potential to serve as a framework for how to use the DO, and other mouse genetic reference populations, to complement and inform human genetic studies of complex disease.

## Acknowledgements

Research reported in this publication was supported in part by the National Institute of Arthritis and Musculoskeletal and Skin Diseases of the National Institutes of Health under Award Numbers AR057759 to C.J.R., M.C.H., and C.R.F., and AR077992 to C.R.F. B.A-B was supported in part by a National Institutes of Health, Biomedical Data Sciences Training Grant (5T32LM012416). The authors acknowledge Wenhao Xu (University of Virginia) and the Genetically Engineered Mouse Models (GEMM) core and the University of Virginia Cancer Center Support Grant (CCSG) P30CA044579 from NCI for their support generating *Qsox1* mutant mice. The authors also acknowledge the Yale School of Medicine Department of Orthopaedics and Rehabilitation’s Histology and Histomorphometry Laboratory for all their work. We thank Matt Vincent (The Jackson Laboratory) and Gary Churchill (The Jackson Laboratory) for developing the QTL Viewer software and Neal Magee (University of Virginia) for hosting QTL Viewer on UVA servers. We thank the IMPC for accessibility to BMD data on *Glt8d2* knockout mice (www.mousephenotype.org). The data used for the analyses described in this manuscript were obtained from the IMPC Portal on 11/5/19. The Genotype-Tissue Expression (GTEx) Project was supported by the Common Fund of the Office of the Director of the National Institutes of Health, and by NCI, NHGRI, NHLBI, NIDA, NIMH, and NINDS. The data used for the analyses described in this manuscript were obtained from the GTEx Portal on 01/15/18.

## Ethics declaration

The animal protocol for the characterization of Diversity Outbred mice and the generation and characterization of *Qsox1* mutant mice was approved by the Institutional Animal Care and Use Committee (IACUC) at the University of Virginia.

## METHODS

### Diversity Outbred mouse population and tissue harvesting

A total of 619 (315 males, 304 females) Diversity Outbred (J:DO, JAX stock #0039376) mice, across 11 generations (gens. 23-33) were procured from The Jackson Laboratory at 4 weeks of age. DO mice were fed standard chow (Envigo Teklad LM-485 irradiated mouse/rat sterilizable diet. Product # 7912), and were injected with calcein (30 mg/g body weight) both 7 days and 1 day prior to sacrifice. Mice were weighed and fasted overnight prior to sacrifice. Mice were sacrificed at approximately 12 weeks of age (median: 86 days, range: 76-94 days). Immediately prior to sacrifice, mice were anesthetized with isoflurane, nose-anus length was recorded and blood collected via submandibular bleeding. At sacrifice, femoral morphology (length and width) was measured with digital calipers (Mitoyuto American, Aurora, IL). Right femora were wrapped in PBS soaked gauze and stored in PBS at −20°C. Right tibiae were stored in 70% EtOH at room temperature. Left femora were flushed of bone marrow (which was snap frozen and stored in liquid nitrogen, see below – Single cell RNA-seq of bone marrow stromal cells exposed to osteogenic differentiation media *in vitro*) and were immediately homogenized in Trizol. Homogenates were stored at −80°C. Left tibiae were stored in 10% neutral buffered formalin at 4°C. Tail clips were collected and stored at −80°C.

### Measurement of trabecular and cortical microarchitecture

Right femora were scanned using a 10 μm isotropic voxel size on a desktop μCT40 (Scanco Medical AG, Brüttisellen, Switzerland), following the Journal of Bone and Mineral Research guidelines for assessment of bone microstructure in rodents ^72^. Trabecular bone architecture was analyzed in the endocortical region of the distal metaphysis. Variables computed for trabecular bone regions include: bone volume, BV/TV, trabecular number, thickness, separation, connectivity density and the structure model index, a measure of the plate versus rod-like nature of trabecular architecture. For cortical bone at the femoral midshaft, total cross-sectional area, cortical bone area, medullary area, cortical thickness, cortical porosity and area moments of inertia about principal axes were computed.

### Biomechanical testing

The right femur from each mouse was loaded to failure in four-point bending in the anterior to posterior direction, such that the posterior quadrant is subjected to tensile loads. The widths of the lower and upper supports of the four-point bending apparatus are 7 mm and 3 mm, respectively. Tests were conducted with a deflection rate of 0.05 mm/s using a servohydraulic materials test system (Instron Corp., Norwood, MA). The load and mid-span deflection were acquired directly at a sampling frequency of 200 Hz. Load-deflection curves were analyzed for strength (maximum load), stiffness (the slope of the initial portion of the curve), post-yield deflection, and total work. Post-yield deflection, which is a measure of ductility, is defined as the deflection at failure minus the deflection at yield. Yield is defined as a 10% reduction of stiffness relative to the initial (tangent) stiffness. Work, which is a measure of toughness, is defined as the area under the load-deflection curve. Femora were tested at room temperature and kept moist with phosphate buffered saline during all tests.

### Assessment of bone marrow adipose tissue (MAT)

Fixed right tibiae, dissected free of soft tissues, were decalcified in EDTA for 20 days, changing the EDTA every 3-4 days and stained for lipid using a 1:1 mixture of 2% aqueous osmium tetroxide (OsO_4_) and 5% potassium dichromate. Decalcified bones were imaged using μCT performed in water with energy of 55 kVp, an integration time of 500 ms, and a maximum isometric voxel size of 10 μm (the “high” resolution setting with a 20mm sample holder) using a µCT35 (Scanco). To determine the position of the MAT within the medullary canal and to determine its change in volume, the bone was overlaid. MAT was recorded in 4 dimensions.

### Histomorphometry

Fixed right tibiae were sequentially dehydrated and infiltrated in graded steps with methyl methacrylate. Blocks were faced and 5 μm non-decalcified sections cut and stained with toludine blue to observe gross histology. This staining allows for the observation of osteoblast and osteoclast numbers, amount of unmineralized osteoid and the presence of mineralized bone. Histomorphometric parameters were analyzed on a computerized tablet using Osteomeasure software (Osteometrics, Atlanta, GA). Histomorphometric measurements were made on a fixed region just below the growth plate corresponding to the primary spongiosa.

### Bulk RNA isolation, sequencing and quantification

We isolated RNA from a randomly chosen subset (n=192, 96/sex) of the available mice at the time (mice number 1-417), constrained to have an equal number of male and female mice. Total RNA was isolated from marrow-depleted homogenates of the left femora, using the mirVana™ miRNA Isolation Kit (Life Technologies, Carlsbad, CA). Total RNA-Seq libraries were constructed using Illumina TruSeq Stranded Total RNA HT sample prep kits. Samples were sequenced to an average of 39 million 2 x 75 bp paired-end reads (total RNA-seq) on an Illumina NextSeq500 sequencer in the University of Virginia Center for Public Health Genomics Genome Sciences Laboratory (GSL). A custom bioinformatics pipeline was used to quantify RNA-seq data. Briefly, RNA-seq FASTQ files were quality controlled using FASTQC ^73^, aligned to the mm10 genome assembly with HISAT2 ^74^, and quantified with Stringtie ^75^. Read count information was then extracted with a Python script provided by the Stringtie website (prepDE.py). Finally, we filtered our gene set to include genes that had more than 6 reads, and more than 0.1 transcripts per million (TPM), in more than 38 samples (20% of all samples). This filtration resulted in 23,648 “genes” remaining from an initial set of 53,801 “genes”. (Note that most of these “genes” were defined by StringTie internally as “genes”, but indicate loci – contiguous regions on the genome where the exons of transcripts overlap). Sequencing data is available on GEO (GSE152708).

### Bulk RNA differential expression analyses

RNA-seq data were subjected to a variance stabilizing transformation using the DESeq2 R package^76^, and the 500 most variable genes were used to calculate the principal components using the “PCA” function from the FactoMineR R package^77^. For visualization, age was binarized into “high” and “low”, with “low” defined as age equal to, or less than, the median age at sacrifice (85 days) and “high” defined as age higher than 85 days. Differential expression was then performed using DESeq2, for both sex and bone strength (max load). For differential expression based on sex, we used a design formula of ∼batch+age+sex. For bone strength, we binarized bone strength into “high” and “low” for each sex independently, using the median bone strength value for each sex (35.66 and 37.42 for males and females, respectively). Differential expression was performed using the following design formula: ∼sex+batch+age+bone strength. Log2 fold changes for both differential expression analyses were then shrunken using the lfcShrink() function in DESeq2, using the adaptive t prior shrinkage estimator from the “apeglm” R package^78^.

### Mouse genotyping

DNA was collected from mouse tails from all 619 DO mice, using the PureLink Genomic DNA mini kit (Invitrogen). DNA was used for genotyping with the GigaMUGA array ^19^ by Neogen Genomics (GeneSeek; Lincoln, NE). Genotyping reports were pre-processed for use with the qtl2 R package ^79 80^, and genotypes were encoded using directions and scripts from (kbroman.org/qtl2/pages/prep_do_data.html). Quality control was performed using the Argyle R package ^81^, where samples were filtered to contain no more than 5% no calls and 50% heterozygous calls. Samples that failed QC were re-genotyped. Furthermore, genotyping markers were filtered to contain only tier 1 and tier 2 markers. Markers that did not uniquely map to the genome were also removed. Finally, a qualitative threshold for the maximum number of no calls and a minimum number of homozygous calls was used to filter markers.

We calculated genotype and allele probabilities, as well as kinship matrices using the qtl2 R package. Genotype probabilities were calculated using a hidden Markov model with an assumed genotyping error probability of 0.002, using the Carter-Falconer map function. Genotype probabilities were then reduced to allele probabilities, and allele probabilities were used to calculate kinship matrices, using the “leave one chromosome out” (LOCO) parameter. Kinship matrices were also calculated using the “overall” parameter for heritability calculations.

Further quality control was then performed ^82^, which led to the removal of several hundred more markers that had greater than 5% genotyping errors, after which genotype and allele probabilities and kinship matrices were recalculated. After the aforementioned successive marker filtration, 109,427 markers remained, out of 143,259 initial genotyping markers. As another metric for quality control, we calculated the frequencies of the eight founder genotypes of the DO.

### WGCNA network construction

Gene counts, as obtained above were pruned to remove genes that had fewer than 10 reads in more than 90% of samples. Genes not located on the autosomes or X chromosome were also removed. This led to the retention of 23,335 out of 23,648 genes. Variance-stabilizing transformation (DeSeq2 ^76^) was applied, followed by RNA-seq batch correction using sex and age at sacrifice in days as covariates (sex was not included as a covariate in the sex-specific networks), using ComBat (“sva” R package ^83^). We then used the “WGCNA” R package to generate signed co-expression networks with a soft thresholding power of 4 (power=5 for male networks) ^84, 85^. We used the “blockwiseModules” function to construct networks with a merge cut height of 0.15 and minimum module size of 30. WGCNA networks had 39, 45 and 40 modules for the sex-combined, female and male networks, respectively.

### Bayesian network learning

Bayesian networks for each WGCNA module were learned with the “bnlearn” R package ^86^. Specifically, expression data for genes within a WGCNA module were obtained as above (WGCNA network construction), and these data were used to learn the structure of the underlying Bayesian network using the Max-Min Hill Climbing algorithm (function “mmhc” in bnlearn).

### Construction of the “known bone gene” list

We constructed a list of bone genes using Gene Ontology (GO) terms and the Mouse Genome Informatics (MGI) database ^87, 88^. Using AmiGO2, we downloaded GO terms for “osteo*”, “bone” and “ossif*”, using all three GO domains (cellular component, biological process and molecular function), without consideration of GO evidence codes ^89^. The resulting GO terms were pruned to remove some terms that were not related to bone function or regulation. We then used the MGI Human and Mouse Homology data table to convert human genes to their mouse homologs. We also downloaded human and mouse genes which had the terms “osteoporosis”, “bone mineral density”, “osteoblast”, “osteoclast”, and “osteocyte”, from MGI’s Human – Mouse: Disease Connection (HMDC) database. Human genes were converted to their mouse counterparts as above. GO and MGI derived genes were merged and duplicates were removed. Finally, we removed genes that were not expressed in our dataset. That is to say, they were not considered in generating the WGCNA modules or Bayesian networks.

### Bone Associated Node (BAN) analysis

We used a custom script that utilized the “igraph” R package to perform BAN analysis ^90^. Briefly, within a Bayesian network underlying a WGCNA module, we counted the number of neighbors for each gene, based on a neighborhood step size of 3. Neighborhood sizes also included the gene itself. BANs were defined as genes that were more highly connected to bone genes than would be expected by chance. We merged all genes from all Bayesian networks together in a matrix, and removed genes that were unconnected or only connected to 1 neighbor (neighborhood size <=2). We then pruned all genes whose neighborhood size was greater than 1 standard deviation less than the mean neighborhood size across all modules. These pruning steps resulted in 15,546/17,264, 13,328/16,446 and 13,541/17,402 genes remaining for the full, male and female Bayesian networks, respectively.

Then, for each gene, we calculated if they were more connected to bone genes in our bone list (see construction of bone list above) than expected by chance using the hypergeometric distribution (phyper, R “stats” package). The arguments were as follows: q: (number of genes in neighborhood that are also bone genes) – 1; m: total number of bone genes in our bone gene set; n: (number of genes in networks prior to pruning) – m; k: neighborhood size of the respective gene; lower.tail = false. False discovery rates were calculated using p.adjust().

### GWAS-eQTL colocalization

We converted mouse genes with evidence of being a BAN (P<=0.05) to their human homologs using the MGI homolog data table. If the human homolog was within 1Mbp of a GWAS association, we obtained all eQTL associations within +/- 200 kb of the GWAS association in all 48 tissues of version 7 of the Gene-Tissue Expression Project (GTEx). These eQTL variants were colocalized with the GWAS variants, using the R “coloc” package, using the coloc.abf() function ^91^. This returned posterior probabilities (PP) for five hypotheses:

- H0: No association with either trait.
- H1: Association with trait 1, not with trait 2.
- H2: Association with trait 2, not with trait 1.
- H3: Association with traits 1 and 2, two independent SNPs.
- H4: Association with traits 1 and 2, one shared SNP.

Genes were considered colocalizing if PPH4 >= 0.75.

### Gene ontology

Gene ontology analysis for WGCNA modules was performed for each individual module using the “topGO” package in R ^92^. Enrichment tests were performed for the “Molecular Function”, “Biological Process” and “Cellular Component” ontologies, using all genes in the network. Enrichment was performed using the “classic” algorithm with Fisher’s exact test. P-values were not corrected for multiple testing.

### Assessing the expression of *Glt8d2* and *Sertad4* in publicly available bone cell data

We used bioGPS expression data from GEO (GSE10246) to assay the expression of *Sertad4*, *Glt8d2,* and *Qsox1* in osteoblasts and osteoclasts^50^. We also downloaded the data from GEO (GSE54461) to query expression in primary calvarial osteoblasts.

### Analysis of BMD data on *Glt8d2^-/-^* mice from the IMPC

The International Mouse Knockout Consortium ^60^ and the IMPC^93^ have generated and phenotyped mice harboring null alleles for *Glt8d2* (*Glt8d2^tm1a(KOMP)Wtsi^,Glt8d2^-/-^*) (N=7 females and N=7 males). Phenotypes for the appropriate controls (C57BL/6) were also collected (N=1,466 females and N=1,477 males). A description of the battery of phenotypes collected on mutants can be found at (https://www.mousephenotype.org/impress/PipelineInfo?id=4). The mice were 14 weeks of age at DEXA scanning and both sexes were included. We downloaded raw BMD, body weight and metadata for *Glt8d2* mutants from the IMPC webportal (http://www.mousephenotype.org). These data were analyzed using the PhenStat R package ^94^. PhenStat was developed to analyze data generated by the IMPC in which a large number of wild-type controls are phenotyped across a wide-time range in batches and experimental mutant animals are tested in small groups interspersed among wild-type batches. We used the Mixed Model framework in PhenStat to analyze BMD data. The mixed model framework starts with a full model (with fixed effects of genotype, sex, genotype x sex and weight and batch as a random effect) and ends with final reduced model and genotype effect evaluation procedures ^94, 95^.

### QTL mapping

Phenotypes that notably deviated from normality were log_10_-transformed (the MAT phenotypes as well as PYD and W_py_ were transformed after a constant of 1 was added). Then, QTL mapping with a single-QTL model was performed via a linear mixed model using the “scan1” function of the “qtl2” R package. A kinship matrix as calculated by the “leave one chromosome out” method was included. Mapping covariates were sex, age at sacrifice in days, bodyweight, and DO mouse generation. Peaks were then identified with a minimum LOD score of 4 and a peak drop of 1.5 LODs. To identify significant QTL peaks, we permuted each phenotype scan 1000 times (using the “scan1perm” function of the “qtl2” package) with the same mapping covariates as above, and calculated the significance threshold for each phenotype at a 5% significance level. Heritability for the phenotypes was calculated using the “est_herit” function of the “qtl2” R package, using the same covariates as above, but with a kinship matrix that was calculated using the “overall” argument.

### DO eQTL mapping

Variance stabilizing transformation was applied to gene read counts from above using the “DESeq2” R package, followed by quantile-based inverse Normal transformation^96^. Then, hidden determinants of gene expression were calculated from these transformed counts, using Probabilistic Estimation of Expression Residuals (PEER)^97^. 48 PEER factors were calculated using no intercept or covariates. Sex and the 48 PEER covariates were used as mapping covariates, and eQTL mapping was performed using the “scan1” function, as above. To calculate a LOD score threshold, we randomly chose 50 genes and permuted them 1000 times, as above. Since all genes were transformed to conform to the same distribution, we found that using 50 was sufficient. Thresholds were set as the highest permuted LOD score each for autosomal chromosomes and the X-chromosome (10.89 and 11.55 LODs, respectively). Finally, we identified peaks as above, and defined eQTL as peaks that exceeded the LOD threshold and were no more than 1Mbp away from their respective transcript’s start site, as defined by the Stringtie output.

### Merge analysis

We performed merge analysis, a previously published approach, using the SNP-association methods in the “qtl2” R package ^61, 80^. For each DO mouse QTL or eQTL peak, we imputed all variants within the 95% confidence interval of a peak, and tested each variant for association with the respective trait. This was performed using the “scan1snps” function of the “qtl2” R package, with the same mapping covariates for QTL or eQTL, respectively. Then, we identified “top” variants by taking variants that were within 85% of the maximum SNP association’s LOD score. For conditional analyses using a variant, we performed the same QTL scan as above, but included the genotype of the respective SNP as an additive mapping covariate, encoding it as a 0, 0.5 or 1, for homozygous alternative, heterozygous or homozygous reference, respectively.

### BMD-GWAS overlap

To identify BMD GWAS loci that overlapped with our DO mouse associations, we defined a mouse association locus as the widest confidence interval given all QTL start and end CI positions mapping to each locus. We then used the UCSC liftOver tool ^98^ (minimum ratio of bases that must remap = 0.1, minimum hit size in query = 100000) to convert the loci from mm10 to their syntenic hg19 positions. We then took all genome-wide significant SNPs (P <=5 x 10^-8^) from the Morris *et al.* GWAS for eBMD and the Estrada *et al.* GWAS for FNBMD and LSBMD, and identified variants that overlapped with the syntenic mouse loci (“GenomicRanges” R package ^99^).

### SIFT annotations

SIFT annotations for merge analysis missense variants were queried using Ensembl’s Variant Effect Predictor tool (https://useast.ensembl.org/Tools/VEP) ^100^. All options were left as default.

### Prior ML QTL mapping

The cohorts used for the earlier QTL mapping of ML consisted of 577 Diversity Outbred mice from breeding generations G10 and G11 ^62^. G10 cohort mice consisted of both males and females fed a defined synthetic diet (D10001, Research Diets, New Brunswick, NJ), and were euthanized and analyzed at 12–15 weeks of age. G11 cohort mice were all females fed a defined synthetic diet (D10001, Research Diets, New Brunswick, NJ) until 6 weeks of age, and were then subsequently fed either a high-fat, cholesterol-containing (HFC) diet (20% fat, 1.25% cholesterol, and 0.5% cholic acid) or a low-fat, high protein diet (5% fat and 20.3% protein) (D12109C and D12083101, respectively, Research Diets, New Brunswick, NJ), and were euthanized and analyzed at 24–25 weeks of age. Mice were weighed and then euthanized by CO_2_ asphyxiation followed by cervical dislocation. Carcasses were frozen at −80°C. Subsequently, the femur was dissected and length, AP width, and ML width was measured two independent times to 0.01 mm using digital calipers. Mice were genotyped using the MegaMUGA SNP array (GeneSeek; Lincoln, NE) designed with 77,800 SNP markers, and QTL mapping was performed as above, but with the inclusion of sex, diet, age and weight at sacrifice as additive covariates.

### Generation of *Qsox1* mutant mice

*Qsox1* knockout mice used in this study were generated using the CRISPR/Cas9 genome editing technique essentially as reported in ^101^. *Qsox1* knockout mice used in this study were generated using the CRISPR/Cas9 genome editing technique essentially as reported in ^101^. Briefly, Cas9 enzyme that was injected into B6SJLF2 embryos (described below) was purchased from (PNA Bio) while the guide RNA (sgRNA) was designed and synthesized as follows: the 20 nucleotide (nt) sequence that would be used to generate the sgRNA was chosen using the CRISPR design tool developed by the Zhang lab (crispr.mit.edu). The chosen sequence and its genome map position is homologous to a region in Exon 1 that is ∼225 bp, 3’ of the translation start site and ∼20bp 5’ of the Exon1/Intron1 boundary (**Supplemental Table 21**). To generate the sgRNA that would be used for injections oligonucleotides of the chosen sequence, as well as the reverse complement (**Supplemental Table 21**), primers 1 and 2, respectively), were synthesized such that an additional 4 nts (CACC and AAAC) were added at the 5’ ends of the oligonucleotides for cloning purposes. These oligonucleotides were annealed to each other by combining equal molar amounts, heating to 90°C for 5 min. and allowing the mixture to passively cool to room temperature. The annealed oligonucleotides were combined with BbsI digested pX330 plasmid vector (provided by the Zhang lab through Addgene; https://www.addgene.org/) and T4 DNA ligase (NEB) and subsequently used to transform Stbl3 competent bacteria (Thermo Fisher) following the manufacturer’s’ protocols. Plasmid DNAs from selected clones were sequenced from primer 3 (**Supplemental Table 21**) and DNA that demonstrated accurate sequence and position of the guide were used for all downstream applications. The DNA template used in the synthesis of the sgRNA was the product of a PCR using the verified plasmid DNA and primers 4 and 5 (**Supplemental Table 21**). The sgRNA was synthesized via *in vitro* transcription (IVT) by way of the MAXIscript T7 kit (Thermo Fisher) following the manufacturer’s protocol. sgRNAs were purified and concentrated using the RNeasy Plus Micro kit (Qiagen) following the manufacturer’s protocol.

B6SJLF1 female mice (Jackson Laboratory) were super-ovulated and mated with B6SJLF1 males. The females were sacrificed and the fertilized eggs (B6SJLF2 embryos) were isolated from the oviducts. The fertilized eggs were co-injected with the purified Cas9 enzyme (50 ng/μl) and sgRNA (30 ng/μl) under a Leica inverted microscope equipped with Leitz micromanipulators (Leica Microsystems). Injected eggs were incubated overnight in KSOM-AA medium (Millipore Sigma). Two-cell stage embryos were implanted on the following day into the oviducts of pseudo pregnant ICR female mice (Envigo). Pups were initially screened by PCR of tail DNA using primers 6 and 7 with subsequent sequencing of the resultant product from primer 8, when the PCR products suggested a relatively large deletion had occurred in at least one of the alleles (**Supplemental Table 17**). For those samples which indicated a small or no deletion had occurred PCR of tail DNA using primers 9 and 10 was performed with subsequent sequencing of the resultant products from primer 11 (**Supplemental Table 21**). Finally, deletions were fully characterized by ligating, with T4 DNA ligase (NEB), the PCR products from either primer pairs 6/7 or 9/10 with the plasmid vector pCR 2.1 (Thermo Fisher) followed by transformation of One Shot Top 10 chemically competent cells (Thermo Fisher) following the manufacturers recommendations (**Supplemental Tables 19 and 20**).

The resulting founder mice (see **Supplemental Table 19**) were mated to C57BL/6J mice (Jackson Laboratory), with CRISPR/Cas9-deletion heterozygous F1 offspring from the 1^st^ and 2^nd^ litters mated to generate the F2 offspring used in the study of bone related properties reported herein. In addition, mouse B (**Supplemental Table 19**) was subsequently mated to an SJL/J male (Jackson Laboratory), and the F2 offspring from the heterozygous F1 crosses, as outlined above, were also used in this study. All F1 and F2 mice from all deletion ‘strains’ were genotyped using primer pairs 9/10, with the PCR products sequenced from primer 11 for mice possessing the 7+6 and 1bp deletions (**Supplemental Table 21**). An additional PCR using primers 6 and 7 was performed with tail DNA from mice carrying the 1347 bp and 756 bp deletions; the products from this 2^nd^ PCR assisted in determining between heterozygous and homozygous deleted genotypes (**Supplemental Table 21**).

ML was measured for both femurs using calipers on a population of 12-week old F2 mice and ML was averaged between the two femurs. A linear model with genotype, mutation type, length, and weight was generated separately for males and females. For the sex-combined data, a sex term was also included in the model. ANOVAs were performed using the Anova() function from the “car” R package ^102^. Lsmeans were calculated using the “emmeans” R package ^103^. The same procedure was performed for the AP and FL sex-combined data.

We randomly selected 50 male F2 mice (25 wt + 25 mut) from the same population, and microarchitectural phenotypes were measured as above, but on left femurs. Bone strength was measured as above but in both the AP and ML orientations. A linear model with genotype, mutation type and weight was generated, and lsmeans were calculated using the “emmeans” R package ^103^.

### Measuring Qsox1 activity in serum

Serum was collected via submandibular bleeding from isoflurane anesthetized mice, prior to sacrifice and isolation of femurs for bone trait analysis. Blood samples were incubated at room temperature for 20-30 m followed by centrifugation at 2000 x g for 10 m at 4°C. The supernatants were transferred to fresh tubes and centrifuged again as described above. The 2^nd^ supernatant of each sample was separated into 50-100 µl aliquots, snap frozen on dry ice and stored at −70°C. Only ‘clear’ serum samples were used for determining QSOX1 activity, because pink-red colored samples had slight-moderate activity, presumably due to sulfhydryl oxidase enzymes released from lysed red blood cells.

Sulfhydryl oxidase activity was determined as outlined in Israel *et al.*, 2014 ^104^ with minor modifications. Briefly, serum samples were thawed on wet ice whereupon 5 µl was used in a 200 µl final reaction volume which consisted of 50 mM KPO_4_, pH7.5, 1mM EDTA (both from Sigma), 10 µM Amplex UltraRed (Thermo Fisher), 0.5% (v/v) Tween 80 (Surfact-Amps, low peroxide; Thermo Fisher), 50 nM Horseradish Peroxide (Sigma), and initiated with the addition of dithiothreitol (Sigma) to 50 µM initial concentration. The reactions were monitored with the ‘high-sensitive dsDNA channel’ of a Qubit Fluorimeter (Thermo Fisher) by measuring the fluorescence every 15-30s for 10m. The assay was calibrated by adding varying concentrations (0-3.2 µM) of freshly diluted H_2_O_2_ (Sigma) to the reaction mixture minus serum. Enzyme activity was expressed in units of (pmol H_2_O_2_/min/µl serum) and typically calculated within the first several minutes of the reaction for wild-type and heterozygous mutant mice. It was calculated using the entire 10m of the reaction for homozygous mutant genotypes.

### Single cell RNA-seq of bone marrow stromal cells exposed to osteogenic differentiation media *in vitro*: *Bone marrow isolation*

The left femur was isolated and cleaned thoroughly of all muscle tissue followed by removal of its distal epiphysis. The marrow was exuded by centrifugation at 2000 x g for 30 seconds into a sterile tube containing 35 μl freezing media (90% FBS, 10% DMSO). The marrow was then triturated 6 times on ice after addition of 150 μl ice cold freezing media and again after further addition of 1ml ice cold freezing media until no visible clumps remained prior to being placed into a Mr. Frosty Freezing Container (Thermo Scientific) and stored overnight at −80° C. Samples were transferred the following day to liquid nitrogen for long term storage.

#### Bone marrow culturing

Previously frozen bone marrow samples from 5 DO mice (mouse IDs: 12, 45, 48, 50, and 84) were thawed at 37° C, resuspended into 5 ml bone marrow growth media (Alpha MEM, 10% FBS, 1% Pen/Strep, 0.01% Glutamax), pelleted in a Sorvall tabletop centrifuge at 1000 rpm for 5 minutes at room temperature and then subjected to red blood cell lysis by resuspending and triturating the resultant pellet into 5 ml 0.2% NaCl for 20 seconds, followed by addition and thorough mixing of 1.6% NaCl. Cells were pelleted again, resuspended into 1 ml bone marrow growth media, plated into one well per sample of a 48 well tissue culture plate and placed into a 37° C, 5% CO_2_ incubator undisturbed for 3 days post-plating, at which time the media was aspirated, cells were washed with 1 ml DPBS once and bone marrow growth media was replaced at 300 μl volume. The process was repeated through day 5 post-plating. At day 6 post-plating, cells were washed in same manner; however, we performed a standard *in vitro* osteoblast differentiation protocol, by replacing bone marrow growth media with 300 μl osteogenic differentiation media (Alpha MEM, 10% FBS, 1% Penicillin Streptomycin, 0.01% Glutamax, 50 mg/ml Ascorbic Acid, 1M B-glycerophosphate, 100μM Dexamethasome). Cells undergoing differentiation were assessed for accumulated mineralization on days 4, 6, 8 and 10 of the differentiation process as follows: IRDye 680 BoneTag Optical Probe (Li-Cor Biosciences, product #926-09374) was reconstituted according to the manufacturer’s instructions. On days 3, 5, 7 and 9, 0.006 nmoles were added to each sample. Twenty-four hours later the cells were washed with 0.5 mls DPBS (Gibco, product #14190250) and media was replaced. The cells were then placed on the Odyssey CLx Imaging System (Li-Cor Biosciences) to measure mineralization density as reflected by IRDye 680 BoneTag Optical Probe incorporation. Final values for mineralization were computed by subtracting the average number of fluorescent units recorded in designated background wells from the number of fluorescent units recorded in the sample wells.

#### RNA isolation

The isolation procedure outlined below was inspired by ^105^. Mineralized cultures were washed twice with Dulbecco’s Phosphate Buffered Saline (DPBS). 0.5ml 60mM EDTA (pH 7.4, made in DPBS) was added for 15-minute room temperature (RT) incubation. EDTA solution was aspirated and replaced for a second 15-minute RT incubation. Cultures were then washed with 0.5ml Hank’s Balanced Salt Solution (HBSS) and incubated with 0.5ml 8mg/ml collagenase in HBSS/4mM CaCl_2_ for 10 minutes at 37° C with shaking. Cultures were triturated 10x and incubated for an additional 20 minutes and 37° C. Cultures were then transferred to a 1.5ml Eppendorf tube, and spun at 500 x g for 5 minutes at RT in a Sorvall tabletop centrifuge. Cultures were resuspended in 0.5ml 0.25% trypsin-EDTA (Gibco, Gaithersburg, MD) and incubated for 15 minutes at 37° C. Cultures were then triturated and incubated for an additional 15 minutes. 0.5ml of media were added, triturated and spun at 500 x g for 5 minutes at RT. Cultures were then resuspended in 0.5ml bone marrow differentiation media and cells were counted.

#### Library preparation, sequencing and analysis

The samples were pooled and concentrated to 800 cells/μl in sterile PBS supplemented with 0.1% BSA. The single cell suspension was loaded into a 10x Chromium Controller (10X Genomics, Pleasanton, CA, USA), aiming to capture 8,000 cells, with the Single Cell 3’ v2 reagent kit, according to the manufacturer’s protocol. Following GEM capturing and lysis, cDNA was amplified (13 cycles) and the manufacturer’s protocol was followed to generate sequencing library. The library was sequenced on the Illumina NextSeq500 and the raw sequencing data was processed using CellRanger toolkit (version 2.0.1). The reads were mapped to mm10 mouse reference genome assembly using STAR (version 2.5.1b) ^106^. Overall, 7,188 cells were sequenced, to a mean depth of 57,717 reads per cell. Sequencing data is available on GEO (GSE152806).

Analysis was performed using Seurat ^107, 108^. Features detected in at least 3 cells where at least 200 features were detected were used. We then filtered out cells with less than 800 reads and more than 5800 reads, as well as cells with 10% or more mitochondrial reads. This resulted in 7,092 remaining cells. Expression measurements were multiplied by 10,000 and log normalized, and the 3000 most variable features were identified. The data were then scaled. Cells were then scored by cell cycle markers, and these scores, as well as the percentage of mitochondrial reads, were regressed out ^109^. Finally, clusters were found with a resolution of 1 and the UMAP was generated. An outlier cluster consisting of 13 cells was removed. Cluster cell types were manually annotated after performing differential expression analyses of the expression of genes in each cluster relative to all other clusters (**Supplemental Table 14**), using the Seurat FindAllMarkers() function, with the only.pos=TRUE argument.

## Data availability

Raw genotyping data, calculated genotype and allele probabilities, and R/qtl2 cross files are available from Zenodo at DOI:10.5281/zenodo.4265417. Raw sequencing data is available from the NCBI Gene Expression Omnibus database (GSE152708, GSE152806). Mapped DO mouse QTL and eQTL can be viewed at http://qtlviewer.uvadcos.io/.

GWAS summary statistics used for this study can be downloaded from http://www.gefos.org/?q=content/data-release-2018 and http://www.gefos.org/?q=content/data-release-2012.

## Code availability

Analysis code is available at https://github.com/basel-maher/DO_project

