## Supplemental Figures for "Systems genetics analyses in Diversity Outbred mice inform human bone mineral density GWAS and identify *Qsox1* as a novel determinant of bone strength"

**Supplemental Figure 1. Principal Component Analysis of bulk RNA-seq data. a**) Scree plot showing the percentage of explained variance for the first 10 principal components. **b**) Individuals in PC1 and PC2 space, colored by sex. **c**) Individuals in PC3 and PC4 space, colored by sex. **d**) Individuals in PC1 and PC2 space, colored by batch. **e**) Individuals in PC1 and PC2 space, colored by batch. **f**) Individuals in PC1 and PC2 space, colored by age (binarized, see Methods). **g**) Individuals in PC3 and PC4 space, colored by age (binarized, see Methods).

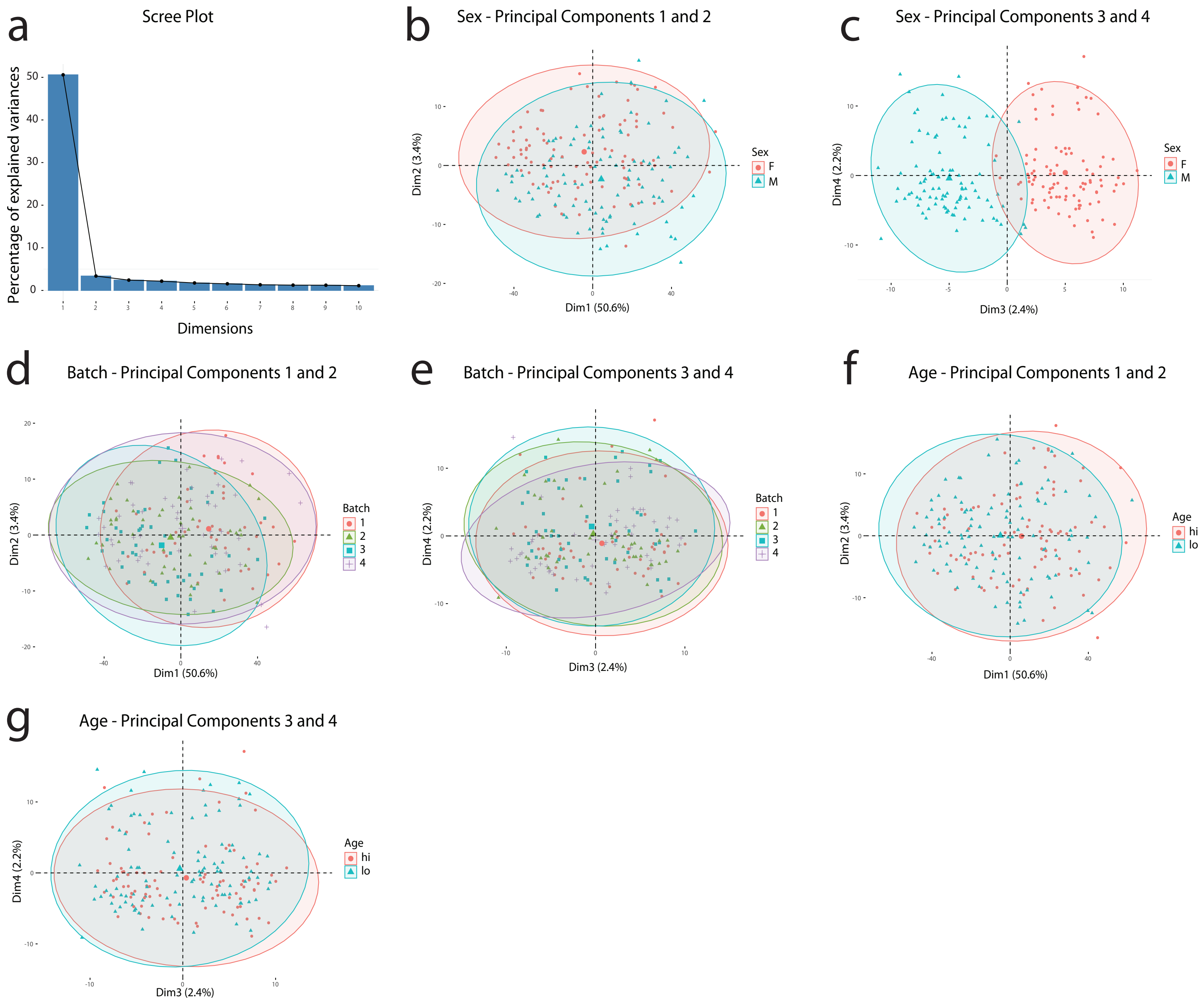

**Supplemental Figure 2. Mineralization of bone marrow-derived stromal cells exposed to osteogenic differentiation media *in vitro*.** During differentiation, cells from each individual DO mouse were assessed for accumulated mineralization by IRDye 680 BoneTag Optical Probe incorporation. The final values for mineralization shown here were computed by subtracting the average number of fluorescent units recorded in designated background wells from the number of fluorescent units recorded in the sample wells. In the cultures from DO mouse #50, there was a much higher percentage of marrow adipogenic lineage precursor cells and a small number of osteoblasts. Consistent with this observation, mouse #50 also demonstrated high levels of marrow adiposity. This is likely the basis of the poor *in vitro* mineralization observed for the cultures from this mouse.

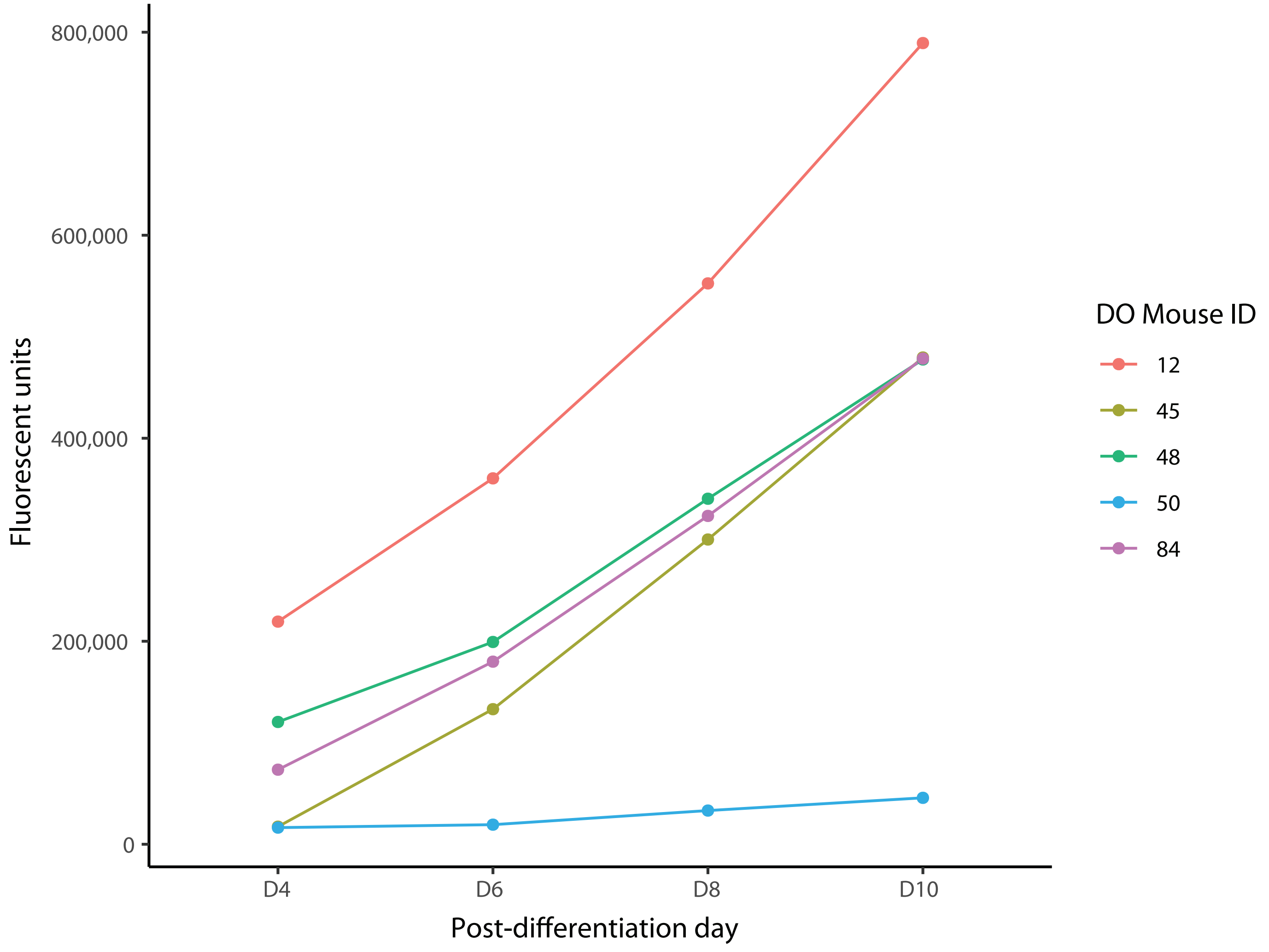

**Supplemental Figure 3. Sex-specific UMAP visulization of single cell RNA-seq expression data on bone marrow stromal cellls cultured in osteogenic differentiation media *in vitro*.** Each point represents a cell.
**a**) *Sertad4* expression in cells from a male DO mouse. **b**) *Glt8d2* expression in cells from a male DO mouse .
**c**) *Sertad4* expression in cells from female DO mince (N=4). **d**) *Glt8d2* expression in cells from female DO mice (N=4).

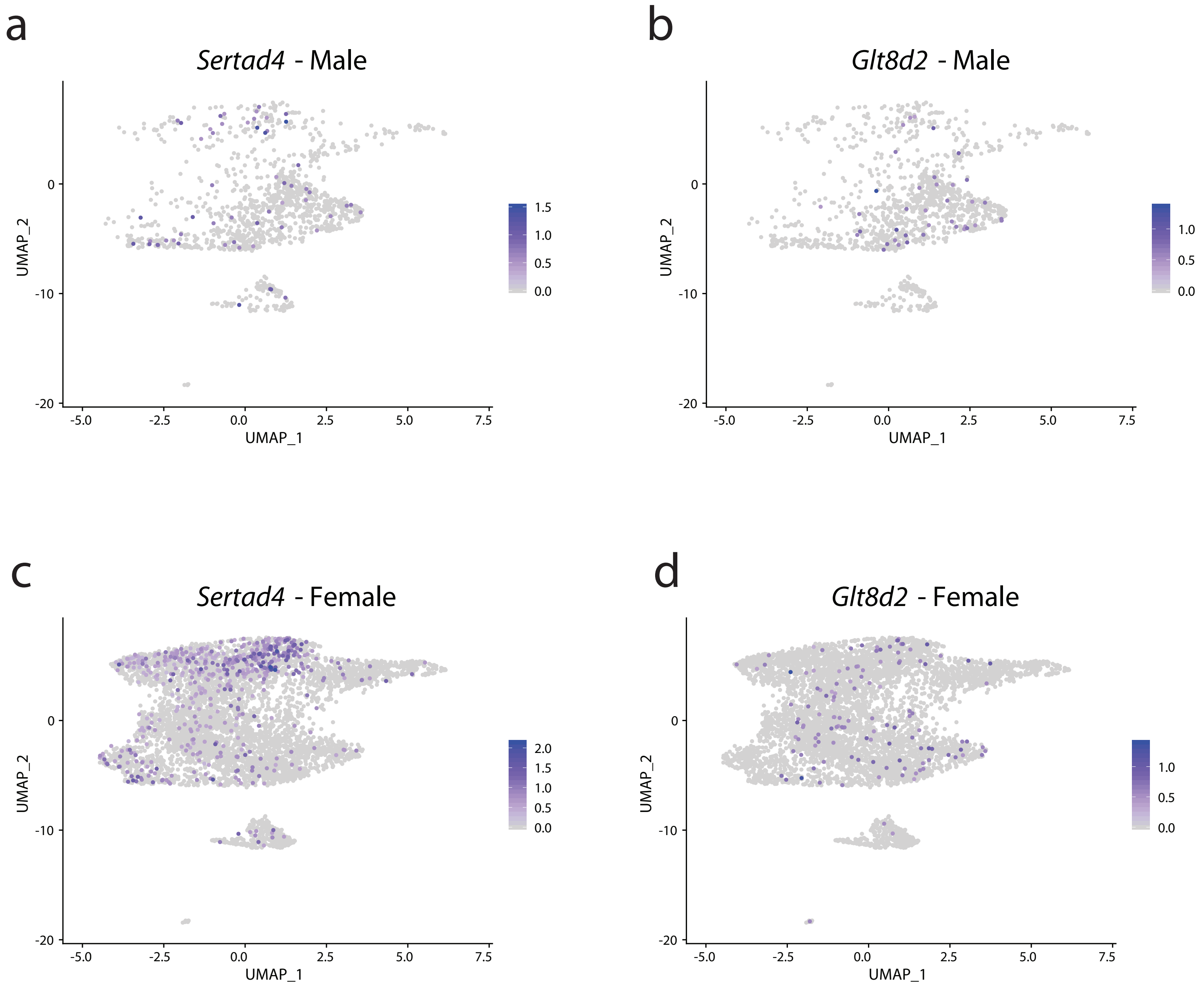

**Supplemental Figure 4. Significant QTL associations.**
Twenty-eight mapped QTL exceeding permutation-based LOD score thresholds (alpha=0.05)

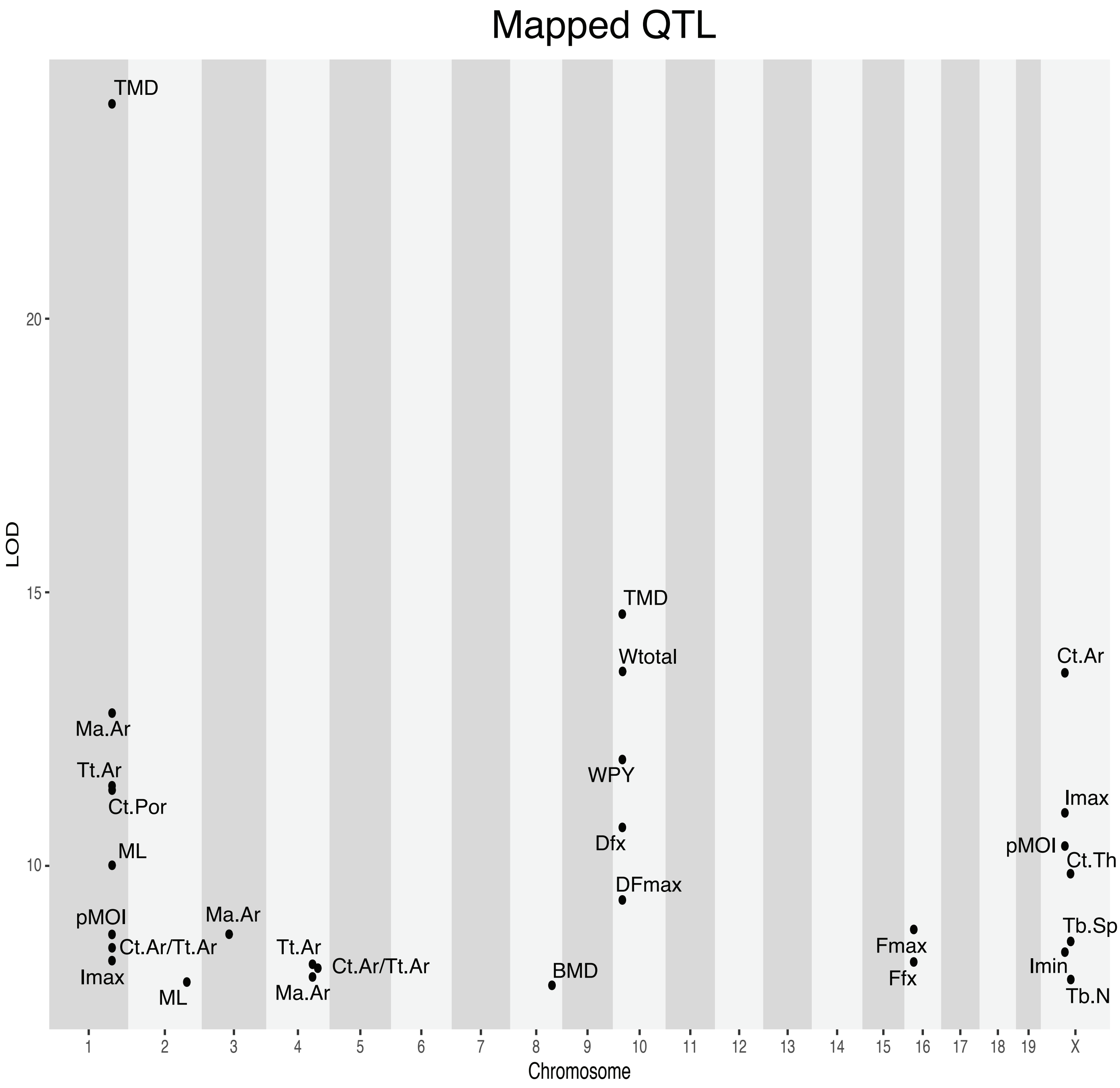

**Supplemental Figure 5. Overlap between BMD GWAS SNPs and QTL loci.** Each panel corresponds to the human syntenic region corresponding to one of the 10 loci identified in the DO. Red circles represent BMD GWAS SNPs (Morris *et al.)* in the locus.
The horizontal lines represent the genome-wide significance threshold (P = $5x10^{-8}$). Not all genes are shown.

a

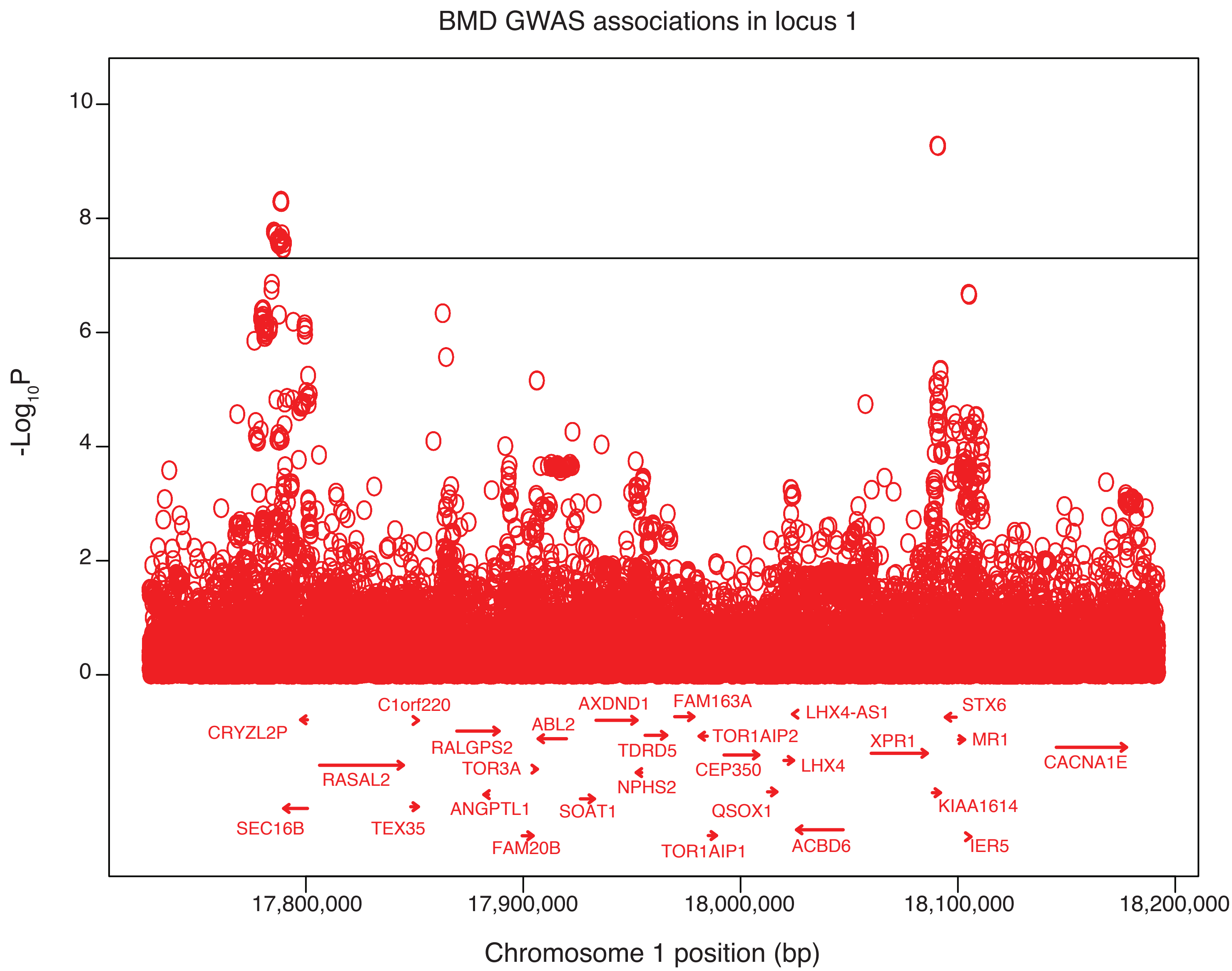

b

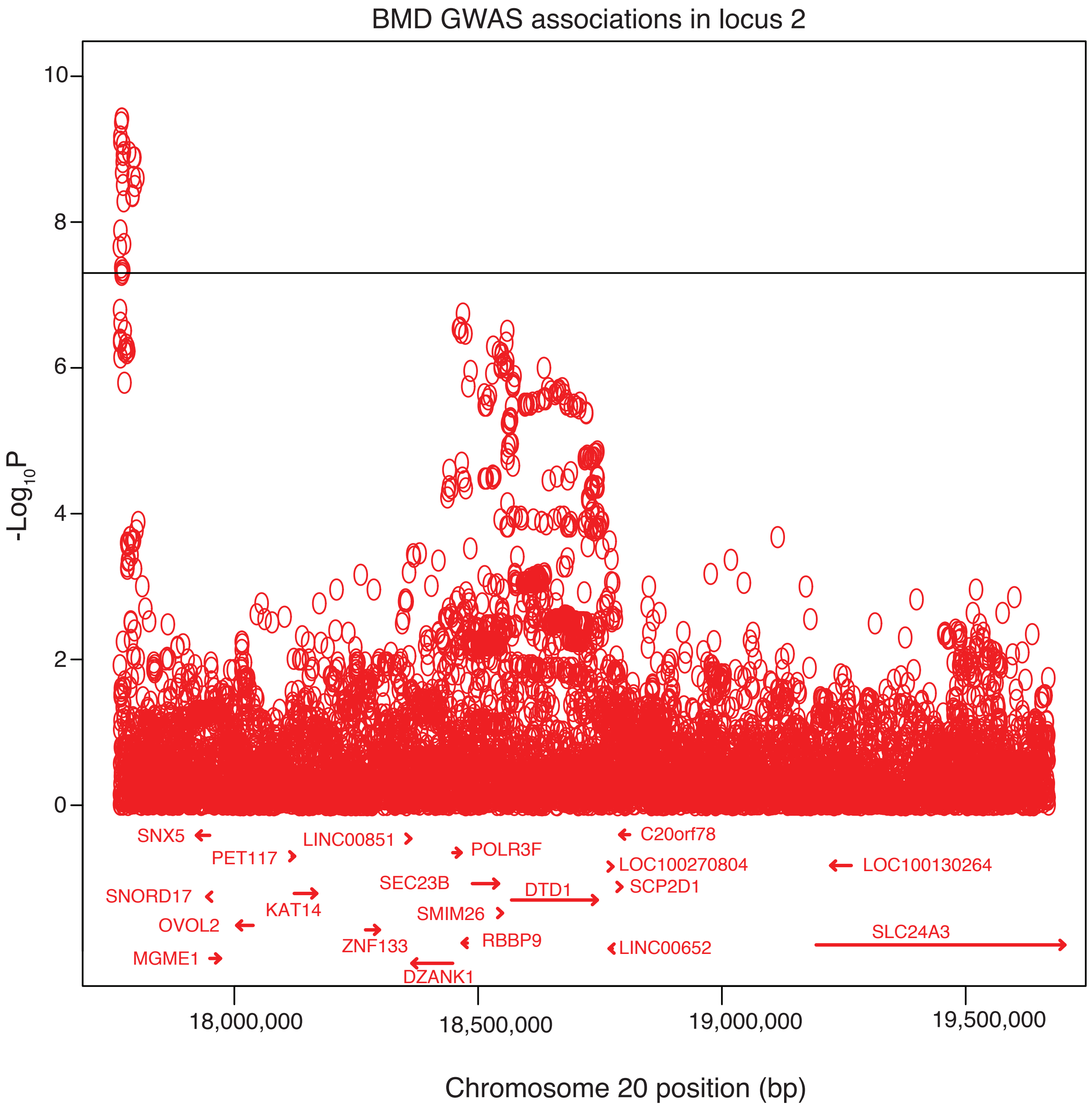

BMD GWAS associations in locus 2
-Log10P
10
8
6
4
2
0
SNX5
PET117
LINC00851
POLR3F
C20orf78
LOC100270804
LOC100130264
SEC23B
DTD1
SCP2D1
SNORD17
KAT14
SMIM26
OVOL2
ZNF133
RBBP9
LINC00652
SLC24A3
MGME1
DZANK1
18,000,000
18,500,000
19,000,000
19,500,000
Chromosome 20 position (bp)

c

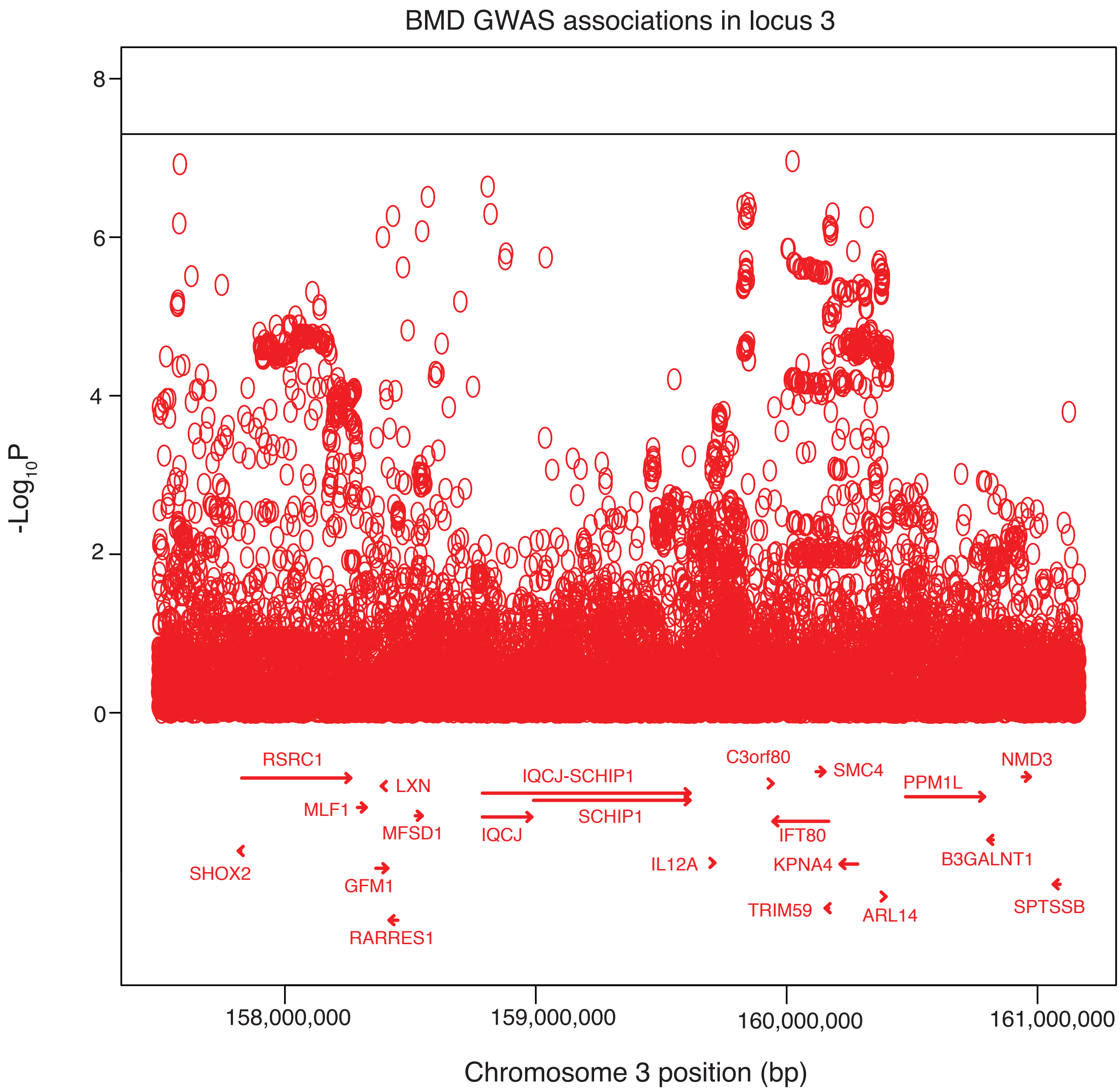
BMD GWAS associations in locus 3
-Log10P
8
6
4
2
0
RSRC1
LXN
MLF1
MFSD1
SHOX2
GFM1
RARRES1
IQCJ-SCHIP1
SCHIP1
IQCJ
C3orf80
SMC4
PPM1L
NMD3
IFT80
IL12A
KPNA4
B3GALNT1
TRIM59
ARL14
SPTSSB
158,000,000
159,000,000
160,000,000
161,000,000
Chromosome 3 position (bp)

d

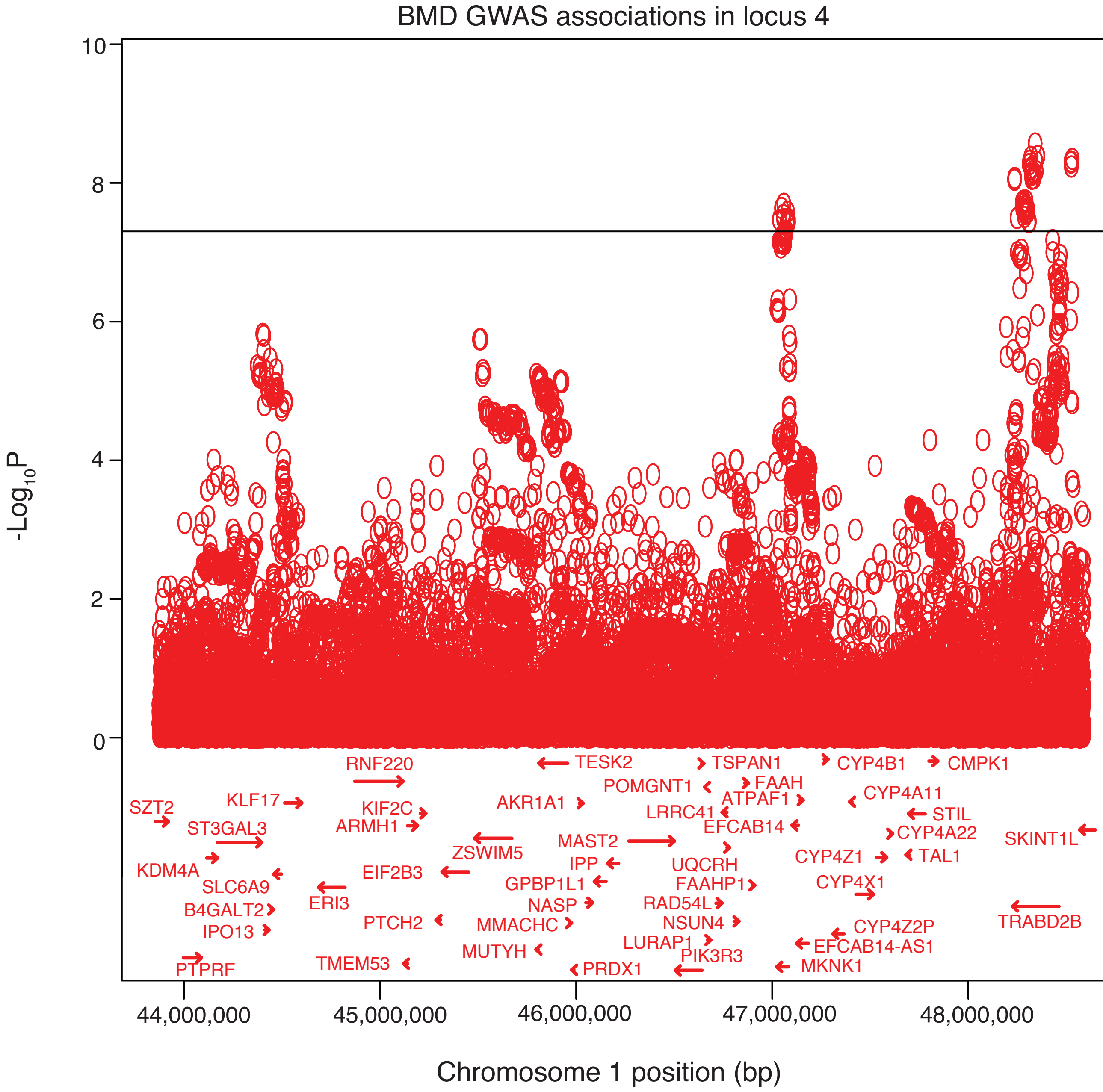
BMD GWAS associations in locus 4
-Log10P
10
8
6
4
2
0
44,000,000
45,000,000
46,000,000
47,000,000
48,000,000
Chromosome 1 position (bp)
RNF220
TESK2
TSPAN1
CYP4B1
CMPK1
POMGNT1
FAAH
SZT2
KLF17
KIF2C
AKR1A1
ATPAF1
CYP4A11
LRRC41
STIL
ST3GAL3
ARMH1
EFCAB14
CYP4A22
SKINT1L
ZSWIM5
MAST2
KDM4A
EIF2B3
IPP
UQCRH
CYP4Z1
TAL1
SLC6A9
GPBP1L1
FAAHP1
CYP4X1
ERI3
NASP
RAD54L
B4GALT2
PTCH2
MMACHC
NSUN4
CYP4Z2P
TRABD2B
IPO13
LURAP1
EFCAB14-AS1
MUTYH
PIK3R3
PTPRF
TMEM53
PRDX1
MKNK1

e

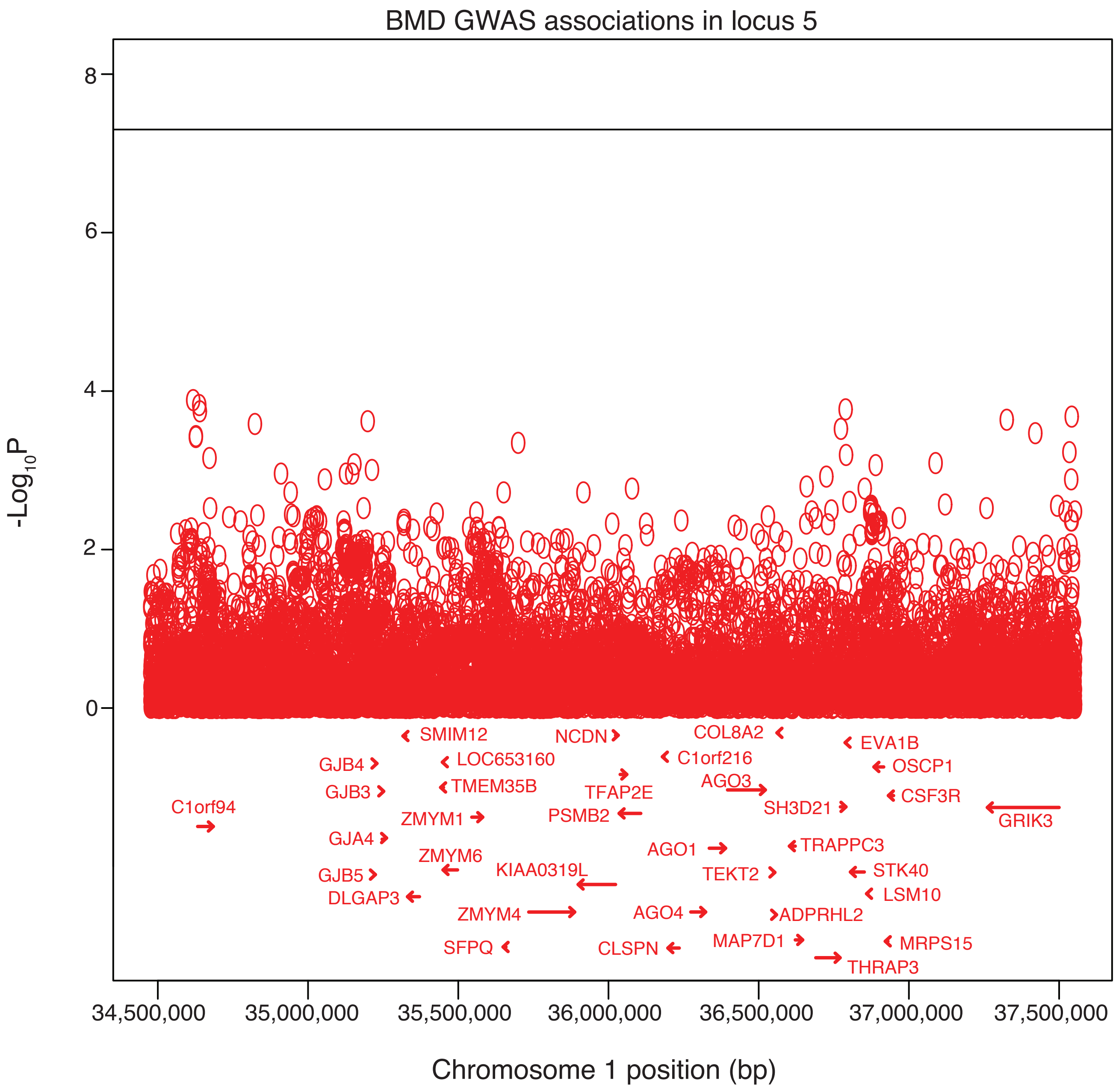

BMD GWAS associations in locus 5
-Log10P
8
6
4
2
0
SMIM12
NCDN
COL8A2
EVA1B
GJB4
LOC653160
C1orf216
OSCP1
GJB3
TMEM35B
TFAP2E
AGO3
CSF3R
C1orf94
ZMYM1
PSMB2
SH3D21
GRIK3
GJA4
AGO1
TRAPPC3
ZMYM6
KIAA0319L
TEKT2
STK40
GJB5
LSM10
DLGAP3
ZMYM4
AGO4
ADPRHL2
MAP7D1
MRPS15
SFPQ
CLSPN
THRAP3
34,500,000
35,000,000
35,500,000
36,000,000
36,500,000
37,000,000
37,500,000
Chromosome 1 position (bp)

f

BMD GWAS associations in locus 6

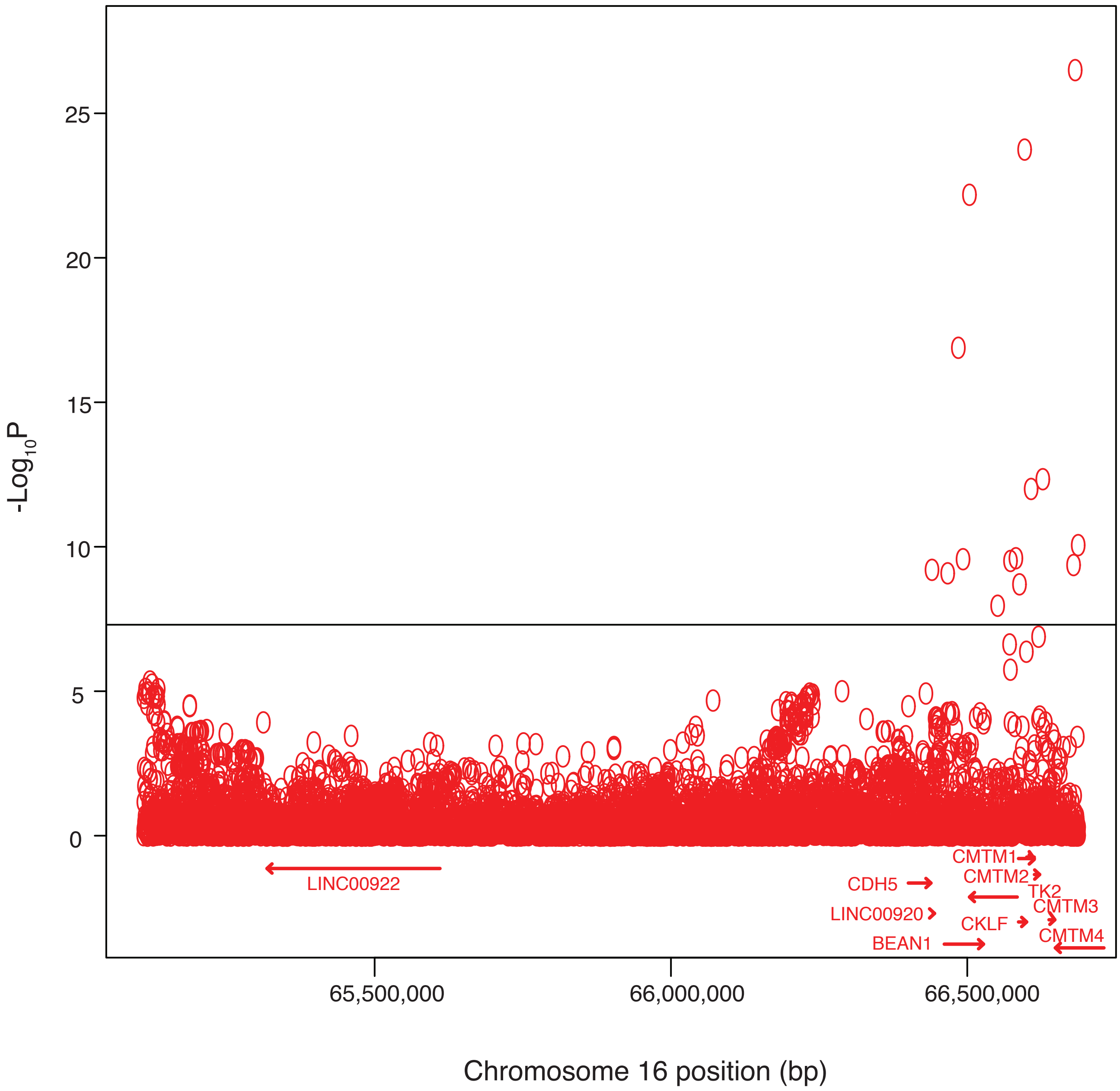

g

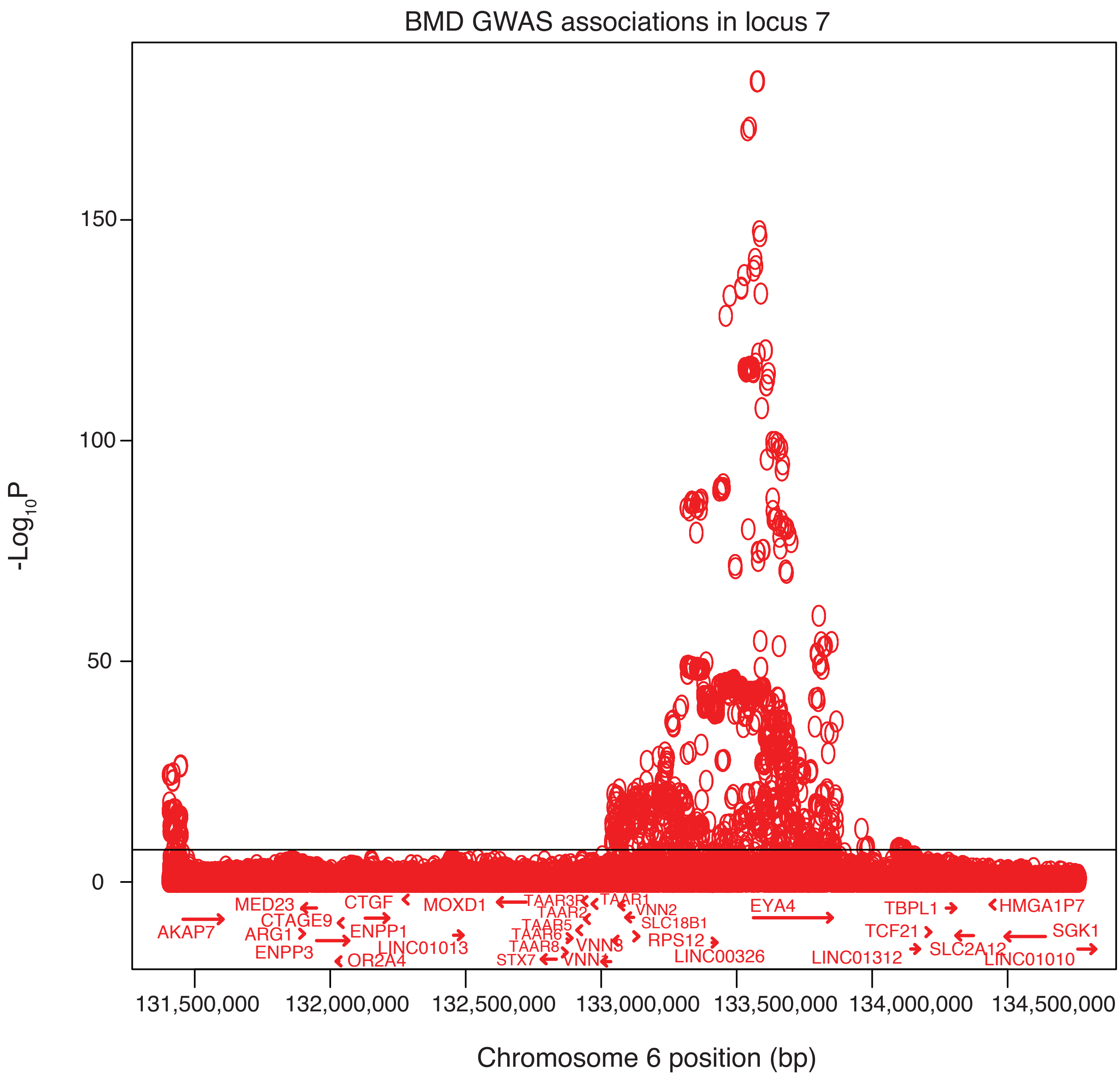

BMD GWAS associations in locus 7
-Log10P
150
100
50
0
MED23
CTGF
MOXD1
TAAR3R
TAAR1
VNN2
EYA4
TBPL1
HMGA1P7
AKAP7
CTAGE9
TAAR2
SLC18B1
ARG1
ENPP1
TAAR5
TCF21
SGK1
ENPP3
LINC01013
TAAR6
RPS12
SLC2A12
TAAR8
VNN3
LINC00326
LINC01312
LINC01010
OR2A4
STX7
VNN1
131,500,000
132,000,000
132,500,000
133,000,000
133,500,000
134,000,000
134,500,000
Chromosome 6 position (bp)

h

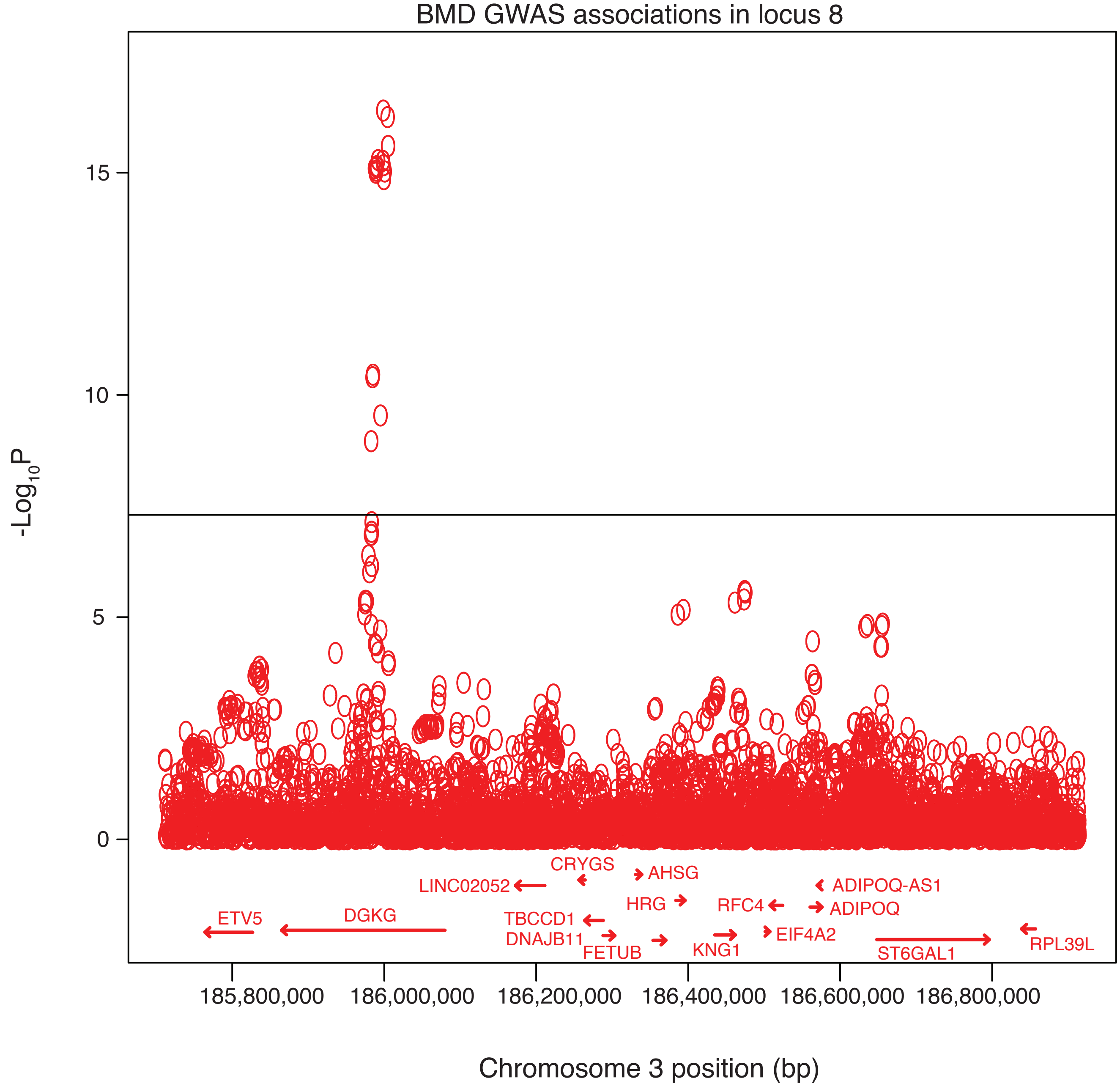
BMD GWAS associations in locus 8
-Log10P
15
10
5
0
CRYGS
AHSG
LINC02052
ADIPOQ-AS1
HRG
RFC4
ADIPOQ
ETV5
DGKG
TBCCD1
EIF4A2
DNAJB11
FETUB
KNG1
ST6GAL1
RPL39L
185,800,000
186,000,000
186,200,000
186,400,000
186,600,000
186,800,000
Chromosome 3 position (bp)

i

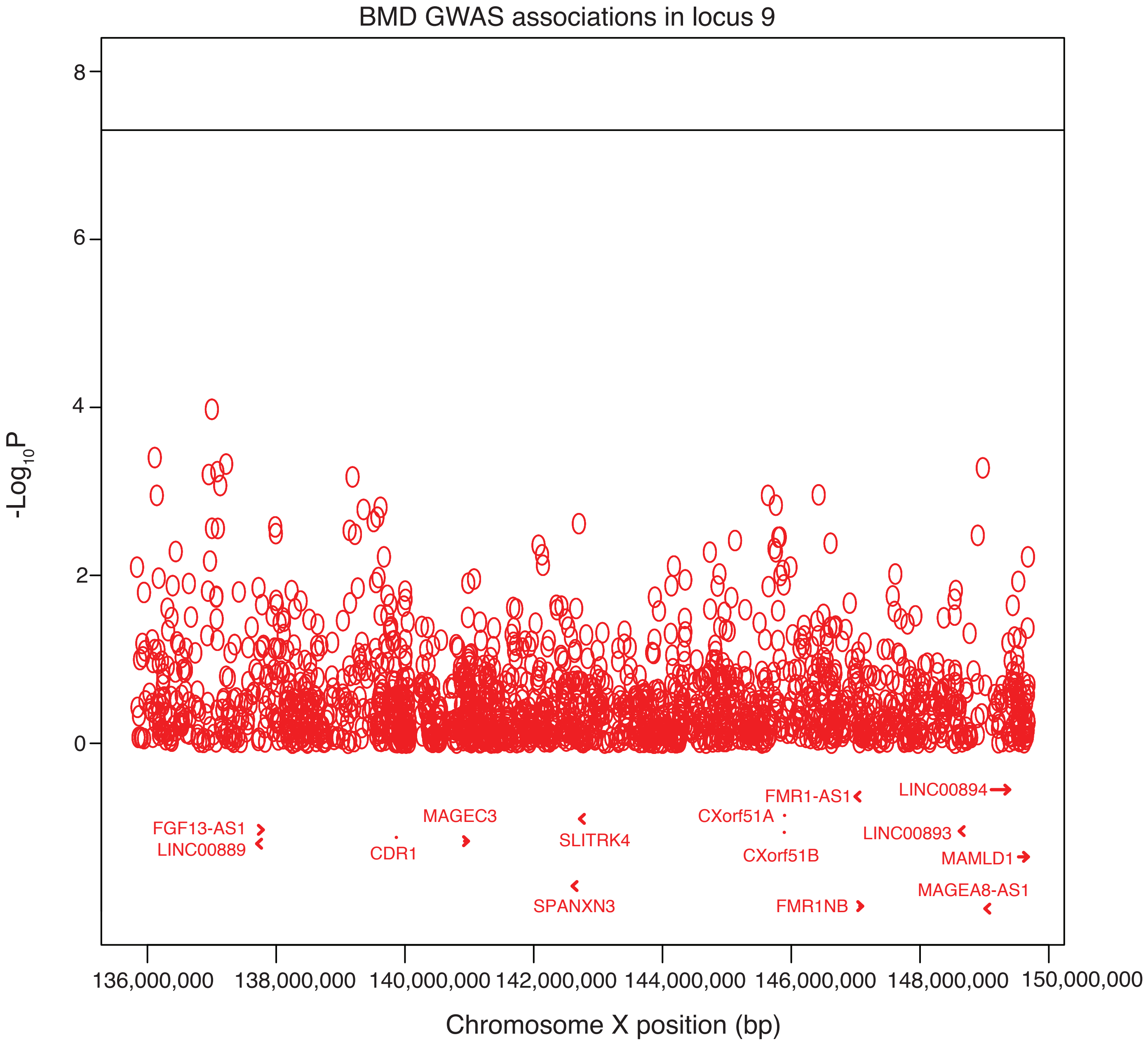
BMD GWAS associations in locus 9
-Log10P
8
6
4
2
0
FGF13-AS1
LINC00889
CDR1
MAGEC3
SLITRK4
SPANXN3
CXorf51A
CXorf51B
FMR1-AS1
FMR1NB
LINC00894
LINC00893
MAMLD1
MAGEA8-AS1
136,000,000
138,000,000
140,000,000
142,000,000
144,000,000
146,000,000
148,000,000
150,000,000
Chromosome X position (bp)

j

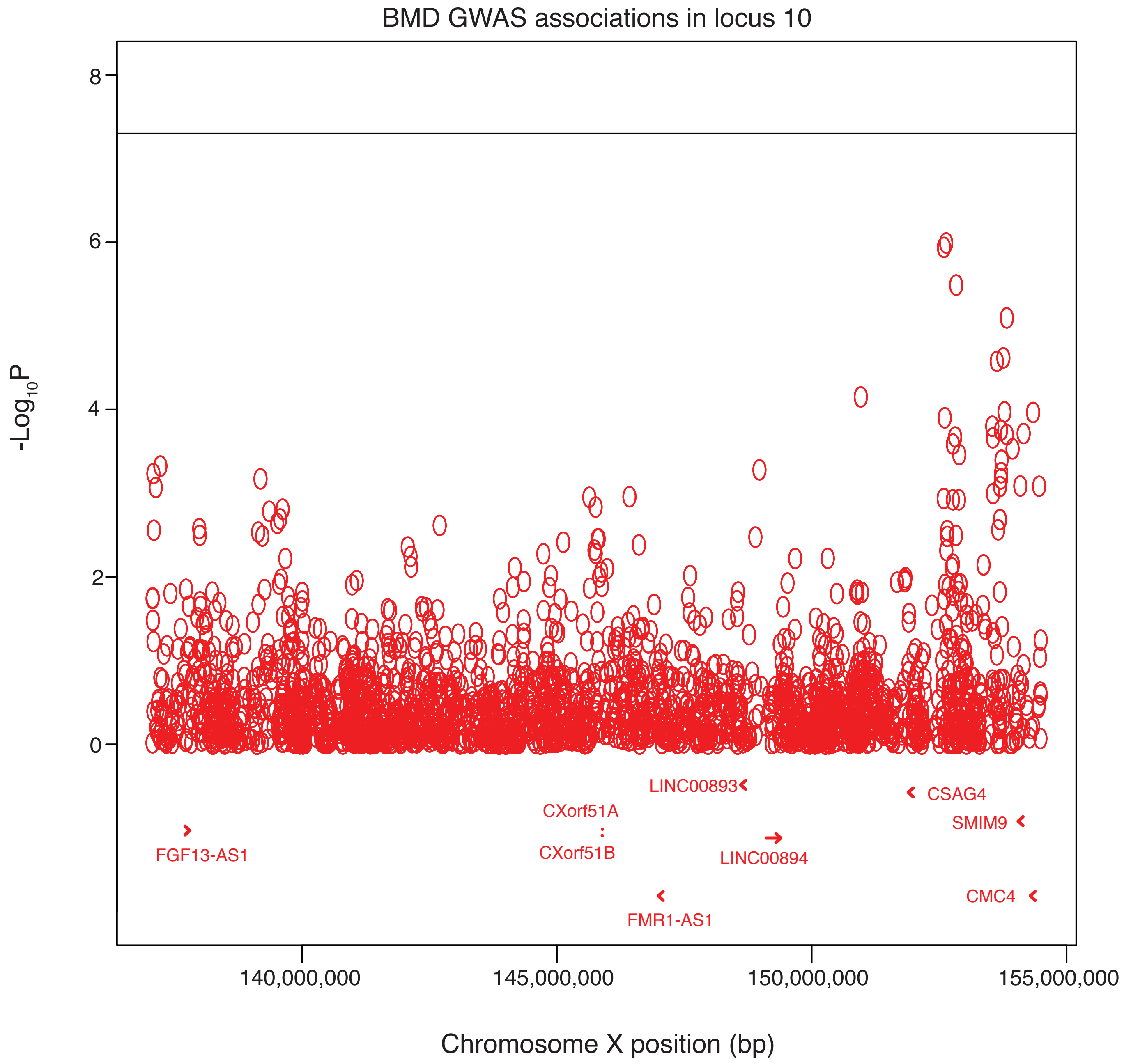

BMD GWAS associations in locus 10
8
6
4
2
0
-Log10P
LINC00893
CSAG4
CXorf51A
SMIM9
FGF13-AS1
CXorf51B
LINC00894
CMC4
FMR1-AS1
140,000,000
145,000,000
150,000,000
155,000,000
Chromosome X position (bp)

**Supplemental Figure 6. ML mapping in a replication cohort.** The top panel shows allele effects for the DO founders for ML in an interval on chromosome 1 (Mbp). Y-axis units are best linear unbiased predictors (BLUPs). The bottom panel shows the QTL scan.

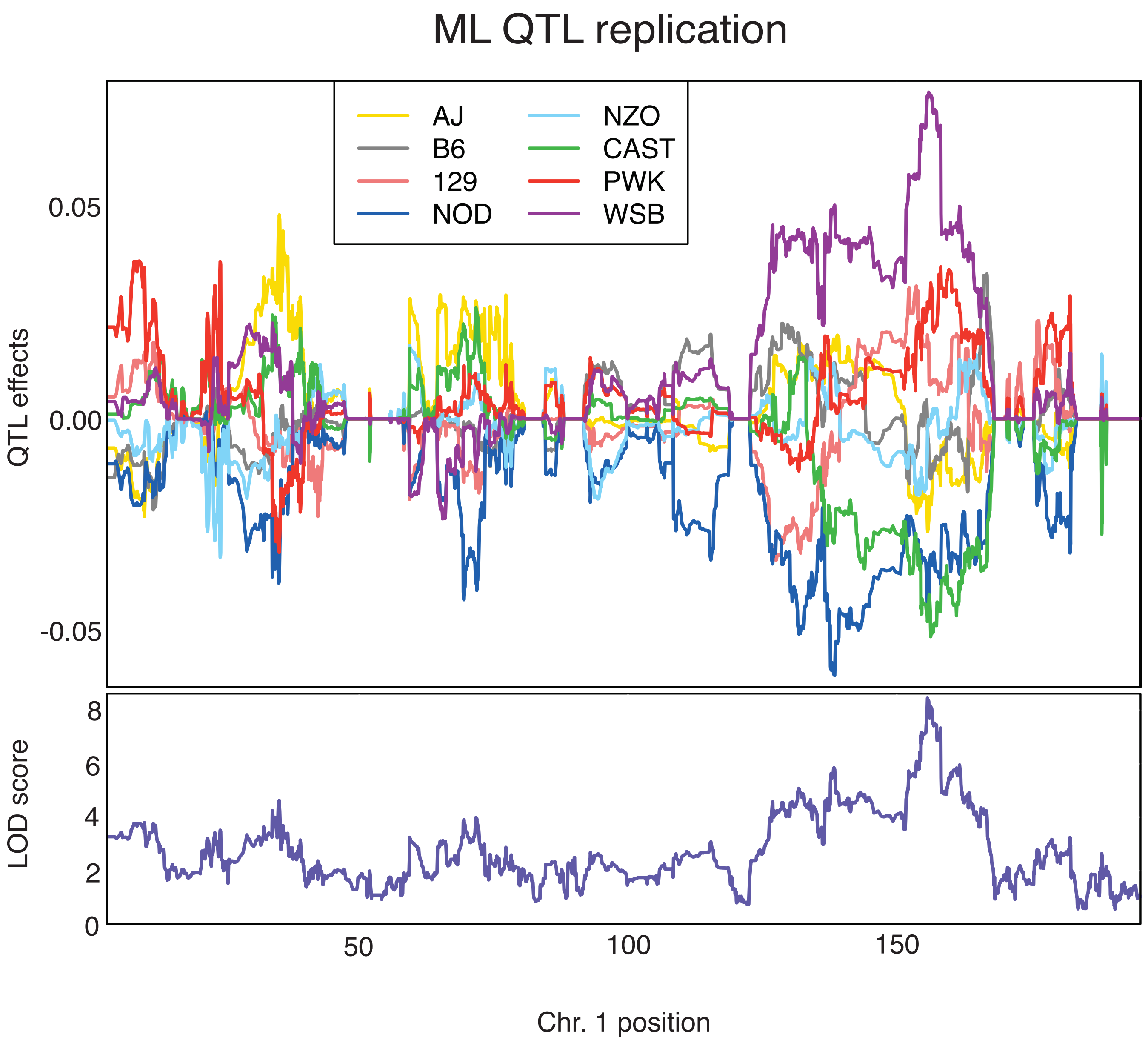
